# Life and death in Trypillia times: Interdisciplinary analyses of the exceptional human remains from the settlement of Kosenivka, Ukraine (3700–3600 BCE)

**DOI:** 10.1101/2023.07.26.550735

**Authors:** Katharina Fuchs, Robert Hofmann, Liudmyla Shatilo, Frank Schlütz, Susanne Storch, Vladislav Chabanyuk, Wiebke Kirleis, Johannes Müller

## Abstract

We present an interdisciplinary analysis of finds from the Trypillia settlement of Kosenivka, Ukraine (ca. 3700–3600 BCE, Trypillia C1), that links information on human, faunal, and botanical remains with archaeological data to provide exceptionally detailed insights into life and death at a Trypillia mega-site. We obtained osteological, palaeopathological, and histotaphonomic data from human bone fragments; performed carbon and nitrogen stable isotopic analysis of human and animal bone to calculate food webs with the software FRUITS; and modelled newly generated radiocarbon dates to refine the site’s chronology. The biological profile of seven identified individuals, some of whom suffered from disease symptoms common in the Chalcolithic, represents a demographic cross-section of the population. The analysis of perimortem cranial trauma suffered by two individuals suggest cases of interpersonal conflict. Food web calculations demonstrate the large contribution of cereals to the protein component of the human diet, which is supported by dental observations, and we suggest that livestock were a major manure producer for crop cultivation.

The most probable scenario for the formation of the Kosenivka find assemblage is a deathly fire event. This makes the site a rare example where the archaeological and osteological results can be used to reconstruct a minimum number of house inhabitants. Following a literature review, we contextualise our analysis by discussing the general lack of human remains from Early and Middle Trypillia sites. The individuals from Kosenivka form part of the less than 0.05% of the total estimated Trypillia population that is represented skeletally; its members were deposited within settlements in the Middle Trypillia stage (until C1), preceding the shift to extramural burials in its late phase (C2).

Our detailed results indicate the huge explanatory potential that has yet to be unlocked in the rare and often poorly preserved bioarchaeological archives of the Cucuteni–Trypillia phenomenon.

## Introduction

Cucuteni−Trypillia societies (CTS) are the earliest Chalcolithic phenomenon in parts of modern-day Romania, Moldavia, and Ukraine ca. 4800–3000 BCE. Succeeding societies with Linear and Bug−Dniester pottery styles, the CTS are the first copper-processing societies, whose settlements later reached sub-urban dimensions, with highly productive agricultural economies [1–3]. Research has focused on site-specific [4–7]; chronological [8,9]; economic [10–14]; and socio-political and cultural aspects [3,15–17], as well as population dynamics [5,18,19]. Compared with the rich archaeological material, human remains are extremely underrepresented due to a general lack of recognisable burial contexts before ca. 3600 BC [20]. Considering the large size of the CTS population groups and the fact that they persisted for approximately 70 generations, the number of finds of human remains is extremely small [5,18,19] and disproportionately concentrated in the western, Cucuteni sphere, rather than the eastern, Trypillia sphere. Thus, we lack not only possible evidence for a *rite de passage* and burial practices, but also human bones as biological archives providing information about the life of the CTS people. This lack of archaeological traces in the form of graves is also seen in many other prehistoric societies [20], and under such circumstances, exceptional preservational conditions are often the only chance to obtain the all-important human bones as archives for reconstructing the lives and deaths of prehistoric populations.

In this paper, we present new results on the exceptional human remains from the Middle Trypillia site of Kosenivka (C1, ca. 3700–3600 BCE). Previously described by Kruts et al. [21] in an archaeological report, the assemblage of unburnt and burnt bones deriving from at least seven individuals not only provides a unique insight into the life and death of the Kosenivka inhabitants, but also revives the discussion on the matter of the “missing dead”. This information is complemented by that from archaeozoological and archaeobotanical finds, which, together with stable isotopic analyses of these finds, allows us to approximate food webs for dietary protein. So far, this kind of isotopic analysis involving human, animal, and plant remains is unique for the CTS.

The aims of this study are twofold. The first is to gain new insights into the life of the Kosenivka individuals through newly obtained bioarchaeological data. To that end, we adopted a multidisciplinary approach that combines archaeological context analysis, osteological, taphonomic, dietary, and palaeopathological aspects; a refined chronological model based on radiocarbon dating; and archaeological analogies. The second is to re-evaluate the Kosenivka bone assemblage regarding the circumstances of the individuals’ death. To that end, we considered analogous human bone assemblages from elsewhere in southeastern and eastern Europe. Our results allowed us to evaluate Kruts et al. [21] assumption that these individuals died in a fire. We found that the results support the scenario of a deadly conflagration.

### Cucuteni–Trypillia phenomena

Sites in the Trypillia area of the Cucuteni–Trypillia phenomenon form a cultural network of agrarian communities characterised by both local and unifying attributes.

In the early phase (Trypillia A1–B1, ca. 4800–4300/4100 BC), the population lived in small and medium-sized settlements of less than 30 ha. In the middle phase (Trypillia B1–B2, B2, and C1, 4300/4100–3600 BC), the population started to aggregate and some huge settlements, measuring 30–320 ha, were established [5,9], particularly in the eastern distribution area, where mega-sites of more than 100 ha are known [2,3]. From its middle phase onwards, the unifying attributes are very specific, including settlement layouts of several concentric circles, some with monumental assembly houses, and a pottery industry producing polychrome painted vessels and various kinds of figurines. Researchers have argued that some of these settlements were pre-urban centres and that they may have had up to ca. 10,000 inhabitants [10,22–25]. In the late phase (C2, ca. 3600–3300/3000 BCE), the domestic sites became smaller once more [23]. The Trypillia mega-site phenomenon thus had a duration of about 300-350 years [26]. Since most of the Trypillia houses are burnt, it is assumed that they were deliberately destroyed when people left the house or settlement [27]. Palaeoecological and palaeodemographic models indicate that the carrying capacity of the local environment’s predominantly forest-steppe areas was never reached [5,11]. The CTS developed sustainable subsistence strategies [13,14], which contributed to the formation of the modern, highly productive chernozem soils and reflect early Anthropocene impact on the landscape [28]. In this regard, we note that the change in settlement pattern from agglomerated back to dispersed, around 3600 BC, occurred for socio-political rather than environmental reasons, specifically social management problems within the mega-sites [16,17].

Because published bioarchaeological information on CTS human remains is very sparse, little is known about the demographic composition and living conditions of CTS populations. The interdisciplinary examination of the individuals from Verteba Cave, located ca. 350 km away, in southwestern Ukraine, near the Siret River, is an important exception, as it offers insights into dietary habits, interpersonal violence and rituals [29]. Palaeogenetic evidence from the 8 individuals from that site [30–32], the single individual from Kolomiytsiv Yar Tract [33], and the four individuals from Moldova [19] illustrates gradual, long-term mixing among main-component Neolithic farmer, European hunter-gatherer, and (proto-)steppe ancestry. One could say that the integrative character of the Trypillia societies became genetically detectable.

### The Middle Trypillia settlement of Kosenivka

In accordance with the local archaeological tradition, the Middle and Late Trypillia phenomenon (stages C1–C2) Kosenovka group is defined by the typology of the cultural material. It includes ca. 25 sites distributed within the interfluve of the Southern Bug and Dnieper rivers [9,21], typically characterised by small settlements (only two exceed 30 ha), ‘surface-type’ adobe houses with wood and daub architecture, table ware with monochrome painting, and characteristic anthropomorphic and zoomorphic figurines [9]. Environmentally, the region was characterised by a forest-steppe landscape [34]. The settlement site of Kosenivka, which has an area of about 80 ha, is located on a plateau north of the Kolodichna River valley and southwest of the modern-day village of Kosenivka, in the Cherkasy region, Umans’kyi district (Fig 1.A). It has been known about since the 1920s. Surveys and archaeological excavations in the years 1982–1988 and 2004 investigated parts of the settlement, including several pits and the burnt remains of six houses. This site, for which the local Kosenovka group was named, is considered to be one of the last Trypillia mega-sites.

**Fig 1.**
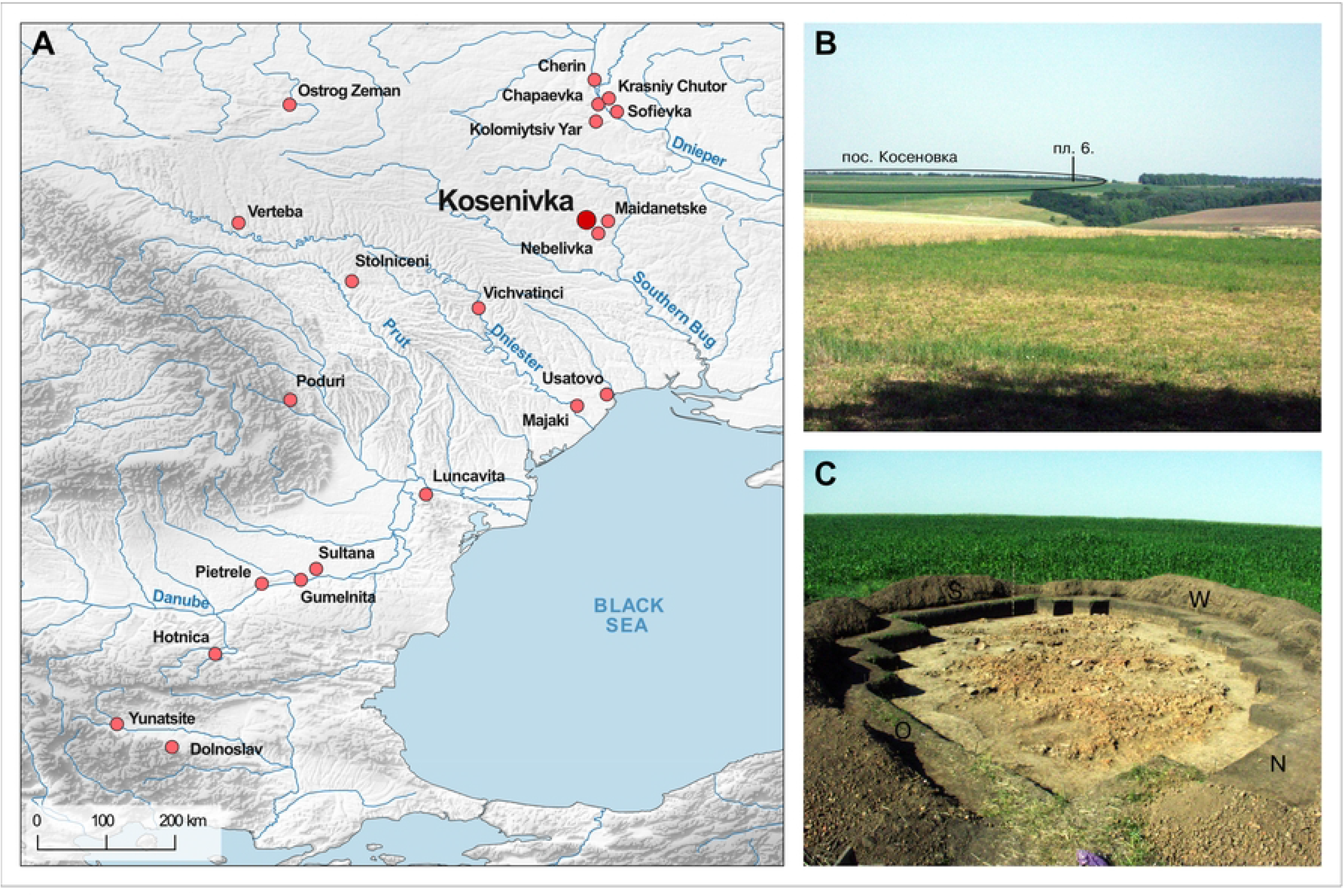
Archaeological context of Kosenivka. (A) Map showing the location of the settlement of Kosenivka and the Chalcolithic sites referred to in the text. (B) Photo showing the location of house 6 within the landscape. (C) Photo showing house 6 being excavated, in 2004 (Map: R. Hofmann. Photos: V. Chabanyuk).

The 2004 excavations by Kruts et al. [21], conducted in 4 m^2^ units (designated with Cyrillic alphabet letters A–E and numbers 1–6), uncovered a chaotic, rectangular accumulation of burnt daub measuring 12 × 4.5 m, which was designated house 6 (Fig 1.B-C). Because it extended only 35 cm below the present-day land surface, the feature was heavily disturbed by modern plough activity. The house is thought to have been covered by more soil originally, which has eroded over time. The northern portion of the house was also disturbed and partly destroyed by a robbery pit. Thus, unfortunately, the architecture of this building is not entirely clear. The discoverers highlighted the absence of the massive platform that is typical for buildings of the adjacent Tomashovka local group. Although the house debris of house 6 suggests that the architecture was much less massive, Kruts et al. [21] assumed it was an elevated or two-story building, with a lower floor and an upper, residential floor. However, due to the damage to the original construction, it is not possible to clearly determine the functions of these two levels. Only three clay installations, which were built directly on the ground, could be clearly associated with the lower floor, including an installation in the northeastern part of the house. One installation could be clearly situated on the upper floor. This linear structure represented the remnant of a partition wall or the southwestern part of the perimeter of an otherwise destroyed oven base.

Larger remains of clay elements with different textures and degrees of combustion in the central interior of the complex are suggestive of spatial divisions that may have included a kiln in the southwestern area, as well as workshops. The find locations of the remains of kitchen ware vessels; granite grinding stones; anthropomorphic and zoomorphic figurines; and a few stone, horn, and bone tools suggest that domestic activities took place inside and outside the building. Faunal bones were found scattered mainly outside of the house.

A total of ca. 80 burnt and 10 unburnt fragments of human bone were found in eight of the 4 m^2^ units, and two further unburnt fragments were found as isolated finds in the overburden. Kruts et al. [21] estimate on the basis of osteological analysis that a minimum number of six individuals are represented—one child, three younger females, and one mature male—but point out that their estimates are uncertain due to the fragmentary condition of the bone.

Calcined bones of three individuals were found within the context of the house; the remains of two of these were found in the area of the clay structure of the assumed oven or workplace in the living room. Unburnt bones of an additional two individuals were found outside of the central accumulation of daub. The researchers state that “If the latter [unburnt bones] can be associated with later burials, the burnt skeletons may be associated with this dwelling. In this case, it must be assumed that the dwelling was burnt by an accidental fire, which killed the people” [21, p. 79, translated by the authors).

In addition to analysing the faunal and human remains from house 6, this study analysed finds of charred cereal grains from house 3 (square 5), excavated during the earlier excavations, for radiocarbon and stable isotopes. A total of 17 radiocarbon dates on human and faunal bone collagen from house 6 and one other house were included in two studies dealing with the relative and absolute chronology of the local Middle and Late Trypillia groups [8,9]. Bayesian modelling conducted in these studies assigns the most probability to the periods 3585–3500 cal BC [8] and 3715–3635 cal BC [9] for these *ploshchadki* (burnt remains of house floors constructed of daub and wood), placing the settlement of Kosenivka at the end of C1 or at the transition from C1 to C2. The modelled dates overlap with those on the human remains from Verteba Cave (3800–3600 BC, [36]).

## Materials and methods

The multidisciplinary approach we applied was aimed at allowing us to explore the bioarchaeological potential of the human remains found at Kosenivka, while taking into account both their rarity and their fragmentary status. We thus chose not to exploit the full range of methods available to us (forgoing, e.g., strontium isotopic analysis) and, following advice to refrain from repeated sampling because genetic analyses on six bone samples and one tooth sample (Ancient DNA Laboratory, Kiel University) did not yield sufficient DNA material, we discontinued our attempt at DNA analysis.

### Human osteology

All human bone fragments and teeth underwent macroscopic osteological examination (naked eye) to estimate age-at-death based on bone- and tooth-specific developmental and age-related changes [37–42]. Infant, juvenile, and mature bones were distinguished on the basis of a qualitative evaluation of microscopic features of bone growth and remodelling [43] for at least five individuals (for protocol, see next subsection). This assessment considered (i) the approximate ratio of lamellar bone to osteons as an indicator of premature bone and (ii) the presence of different generations of osteons as an indicator of mature bone [44,45]. For the microscopic methodology see section Fire impact and bioerosion. Poor skeletal element representation per individual precluded application of morphometric approaches, such as that established by Kerley and others [43,46]. Biological sex was estimated where possible based on landmarks on the pelvis, the skull, and the long bones that are indicative for sexual dimorphism [37,47–52]. For estimating which left and right appendicular elements match, as well as for other metrical aspects [53], we measured bone landmarks, using a sliding calliper.

The minimum number of individuals (MNI) was estimated based on a combination of element duplication per body side, markers for age-at-death, and markers for biological sex, as well as *in situ* proximity of recovery within the archaeological context. In our opinion, the “re-individualisation” that we thus arrived at can be considered as highly probable. A summary of the results can be found in S1 Tables.

### Palaeopathology: Disease and trauma

Human remains offer unique insight into the experiences an individual underwent throughout their life, as manifested in the constantly remodelling skeletal body. Markers of habitual activities; overall appearance; and signs of physical stress, disease, and trauma are all valuable in that respect. Once the soft tissues have decayed, this insight is limited to pathological processes that lasted long enough to result in osseous symptoms before death (inflammation, metabolic disorders, general physiological stress, overload of the musculoskeletal system, oral diseases, and dental manipulations). For traumatic injuries, it is possible to evaluate whether they were intravital or perimortem in nature by assessing signs of healing and fracture patterns.

Due to the poor representation of individual skeletal elements per individual and the fire-induced alterations in the Kosenivka assemblage, it is able to provide relatively limited information in this regard (S1 Tables). Thus, observations on bones and teeth represent the minimum of pathologies and traumas the Kosenivka individuals suffered from. But because published analyses on human remains are so rare, this fragmentary insight still adds significantly to the knowledge of Trypillia life.

Analysis of palaeopathologies and indicators for trauma were performed by macroscopic inspection (naked eye, magnifying glass) and using a digital microscope (Keyence Digital Microscope VHX-500) and a sliding calliper. Unless stated otherwise in the results, diagnostic assumptions associated with our observations of osseous and dental changes and associated diagnostic assumptions can be found in standards provided by Ortner [54], Schultz [55], and Weber and colleagues [56]. The focus of the analyses was on entheseal changes indicating physical activity; inflammatory or other changes on bone surfaces and sinuses indicating infection; and oral categories, such as typical and atypical dental wear, periodontal disease, and caries [57]. Causation of traumatic injuries was inferred based on fracture patterns, fracture locality, and signs of fracture healing, following the forensic and clinical observations published by Weber et al. [58] in order to understand their causation.

### Impact of fire and bioerosion

A central concern of the current research is to formulate possible scenarios that led to the combination of unburnt and burnt bones in the recovered assemblage. One obvious scenario is an accidental conflagration, as suggested by Kruts and colleagues [21]. A second scenario, which takes into consideration that cremations are known for the Late Trypillia stage, is cremation, that is, a mortuary practice involving fire. A third scenario is intramural burials that were burnt later, sometime after their initial deposition, perhaps together with the house structures. To confirm or eliminate each of these three scenarios, it is crucial to date house 6 as reliably as possible and to determine in which taphonomic state the human remains were exposed to fire, that is, whether they were “wet” or “dry”, and to use variation in fire-induced bone alterations to infer combustion temperatures and expansion. We therefore compared macroscopic and microscopic bone alterations of skeletal elements with and without traces of fire (S1 Tables). Key pieces of information are changes to the bone structure induced by (i) physical, chemical, and thermal manifestations of fire and (ii) microbial attack targeting the organic part of the bone. We contextualise the results regarding their localisation within the archaeological structure by focusing on both burnt and non-burnt domestic remains.

Wet (also known as fresh or green) bone consists of hydrated organic and inorganic material that corresponds to its original structure. Bones of living organisms and organisms that have recently died, including in defleshed bones, are found in this wet state. Dry (also known as skeletonised) bone is devoid of water, soft tissue, and bone tissue cells, consisting of just collagen and minerals. Bones from human bodies that have been allowed to naturally decompose, which is often characteristic for archaeological skeletal remains, are typically found in this dry state.

During the transition from wet to dry bone, the collagen is attacked by endogenous and exogenous microbes, a process of bone degradation called bioerosion [59]. The microbial removal of proteins dissolves the bioapatite structure, causing characteristic alterations in the mineralised bone matrix that are evident during microscopic analysis (microscopic focal deconstruction, MDF,44,45). In terrestrial environments, bioerosion is caused by fungal and bacterial attack and distinguished based on a tunnel-like (Wedl tunnels), or focal, shape, which can be observed in bone histology thin-sections [61]. Because the abundance and diversity of microbes and their access to human tissues are associated with, among others, different stages of body decomposition, this histotaphonomic method is used to trace treatment of the corpse during funerary processes and burial environment in forensic and archaeological cases [62,63]. Experimental studies agree that microbial attack to the bone may start soon after death [64,65]. Therefore microbial bone alteration is expected to be absent in human remains that were exposed to fire perimortem, and vice versa (66). In contrast to other studies on bioerosion, we based our assessment of whether the exposure was perimortem or sometime later on the absence or presence of microbial degradation rather than on the extent or form of the microbial alterations [67,68]. The identification of Wedl, budded, longitudinal, or lamellate tunnelling is intended to merely trace the spectrum of microbial activity [61] see S2 Appendix Fig 17).

Bone reacts differently to the impact of fire depending on the condition and structure of the bone and on the temperature and duration of combustion. Generally, in bone, combustion results in loss of incorporated water, decomposition of organic tissue, and reorganisation of the bioapatite crystalline structure [69]. Morphological features indicative of combustion are bone colour, texture, strength, fraction patterns, deformation, and shrinkage. These characteristics are the basis not only of studies dealing with funeral-associated cremation, but also of studies trying to disentangle multi-phased treatments of human remains [70,71]. Thus, the terms “burnt” and “cremated” are not synonymous, as the latter refers to intentional use of fire during the funeral process [69]. Depending on its timing and extent, intentional use of fire may cause different morphological features. Bone assemblages from cremations on funeral pyres typically show a high degree of shrinkage and fragmentation. However, this level of fragmentation is due to the extinguishing of the fire and the subsequent collection and perhaps shattering of the burnt remains to fit them in urn vessels [72]. Diagnostic criteria for the combustion of wet bone stem from experimental work undertaken for archaeological and forensic cases [73–75]. The most accepted indicators are heat-induced cracks and burst Haversian canals, resulting from increasing volume pressure that, in turn, results from water rapidly evaporating under high heat [76]. Other indicators, such as warping and thumbnail fractures, are indicative of, but not decisive for, wet bone combustion [74].

Both microbial and fire effects on bone were analysed through macroscopic and microscopic inspections, which included an evaluation of histological preservation according to the Oxford Histological Index (OHI, [61]). Macroscopic morphological features of fire exposure were examined focusing on colour, fragment size, signs of warping, and fraction patterns [78,79]. Microscopic morphological features of fire exposure were examined through inspections of histological thin-sections, 60μm thick, of seven burnt and three unburnt bone fragments. Where feasible, section preparation targeted compact bone, as it displays micromorphological features and microbial damage well. The thin-sections were viewed under transmitting light using an Olympus SZX7 stereo microscope with an Olympus SC50 camera and the Olympus cellSensEntry software, as well as plain and polarising light using a Nikon Eclipse 50i microscope a Jenoptics PROGRES Gryphax camera and associated software. Observations focused on (i) histological preservation, (ii) fire-induced alterations, and (iii) signs of microbial attack.

### Radiocarbon dating

A total of 23 bone samples (9 unburnt and 5 calcined human bone specimens, 9 faunal unburnt bone specimens) from house 6 were sent to the Poznań Radiocarbon Laboratory, Poland, for bone collagen extraction and accelerator mass spectrometry (AMS) measurement of ^14^C, in 2020 and 2021. Sample pretreatment and analytical procedures followed the https://radiocarbon.pl/en/sample-preparation/ <u>institute’s protocols</u> for unburnt and cremated bone samples, respectively [80–82]. Collagen quality was assessed as being sufficient, as the sample had a C/N anatomic ratio higher than 2.7 and a collagen yield higher than 0.5%. Three out of five samples of calcined bone were not suitable for dating. Measurement took place using the Compact Carbon AMS spectrometer [83].

One charred emmer (*T. dicoccum)* grain from house 3 was sent to the Leibniz Laboratory for Radiometric Dating and Stable Isotope Research, Kiel, Germany, for standard AMS measurement. The grain was examined for impurities under the microscope before a suitable sample amount was taken. Sample pretreatment (with 1% HCl, 1% NaOH at 60°C, and again with 1% HCl) and analytical procedures followed the https://www.leibniz.uni-kiel.de/en/ams-14c-lab/reporting-of-results <u>institute’s protocols</u> for charred organic material. Measurement took place using the HVE 3MV Tandetron 4230 AMS type.

For all samples, conventional ^14^C age was corrected following Stuiver and Polach [84]. Calibration of ^14^C age was performed in the programme OxCal, v. 4.4.165 [85], using the INTCAL20 calibration curve [86].

We included five published dates from unidentified faunal bones lacking stratigraphic information, recovered from house 3 and a not-further-specified domestic feature, in our chronological analyses [8]. Thus, altogether, 26 bone samples are available for the dating of Kosenivka: 20 bone samples from house 6, 5 from house 3, and 1 from the unspecified context (Table 3, S3 Tables).

We generated sum calibrations per house sample set by considering possible inter-laboratory discrepancies. We obtained two ^14^C dates from the same lab for the single human frontal bone, which offered valuable information on intra-individual variability. For house 6, we applied Bayesian modelling taking into account the stratigraphic information on one human skull bone, which was described as having been found “on top on the house debris” [21], suggesting a younger stratigraphic age, and by applying the “boundary” function to the human and faunal bones separately. This house is dated based on the assumption that, unlike the human skeletal remains, the scattered faunal bones are waste remains from when the house was being lived in. We also included the dates obtained for house 3 as a separate unit. This allows us to better evaluate whether the houses were being lived in simultaneously and thus obtain a tighter estimation of the temporal range of the settlement of Kosenivka.

We provide multiple and curve plots for both the crude and the modelled results generated by OxCal (S4 Appendix), as well as the original modelling code (S3 Tables).

### Stable C and N isotope analyses: Flora, fauna, and humans

The recovery of the remains of arable plants and of wild as well as domestic livestock together with the human remains makes Kosenivka a significant site for gaining insights into Trypillia subsistence. We use this unique opportunity to investigate the food web by means of a mixing model approach [87,88] and to approximate the isoscape of the Chalcolithic environment. We do so using stable carbon and nitrogen isotopes from charred cereals and from animal as well as human bone collagen.

### Stable C and N isotopes from cereals

Charred cereal grains of einkorn (*Triticum monococcum*) and emmer (*T. dicoccum)* have been recovered from house 3. We note that these cereal grains’ isotopic signatures may have been distorted by manuring, compared with grains grown in natural conditions. For our calculations, we did not adjust the measured δ^13^C of the cereals for charring, because the literature on the effects of firing is not unanimous and, where it finds effects, these are low. However, we did adjust the measured δ^15^N of the cereals, by −0.5‰, to correct for changes during the charring process [89–94]. For the isoscape reconstruction, we reduced the grain δ^13^C by 2‰ to reconstruct the corresponding δ^13^C of the vegetative plant parts [95,96], in order to allow comparison with species of vegetation that was likely grazed or browsed by animals.

The charred cereal grains were prepared for sampling and analysed at the stable isotope lab at the Department of Earth & Environmental Science, KU Leuven, Belgium. Stable isotopes of carbon and nitrogen contents were measured through combustion using an elemental analyser (ThermoFisher, Flash HT/EA or EA 1110) coupled to an isotope ratio mass spectrometer system (ThermoFisher, Delta V Advantage) via a continuous-flow interface (ThermoFisher, Conglo IV) with the standards IAEA600 (−27.77‰ δ^13^C; 1‰ δ^15^N), Leucine (−13.73‰, 1.07‰), and Tuna (−18.72‰, 13.77‰). Standard deviations on the measured isotopic values are typically below 0.1‰. Isotopic δ-values are expressed relative to the international Vienna PeeDee Belemnite for ^13^C and AIR for ^15^N.

### Stable C and N isotopes from faunal and human bone collagen

For the faunal isotopic signatures, we sampled seven bones of the herbivore species aurochs (*Bos primigenius*), cattle (*B. taurus*), and sheep/goat (*Ovis aries/Capra hircus*). Results from a wild omnivore as well as five unidentified species [8] complete the Kosenivka animal isotopic data set. For human signatures, representing the final consumer of a food web, unburnt specimens provided suitable sample material. We used the same samples that we used for the radiocarbon samples, that is, bones from three individuals identified by their cranial remains (individuals 5–7), as well as five isolated bones that could not be associated with one of the numbered individuals. The bone collagen values represent a mixed isotopic signal of the food consumed by the individuals or animals during the last years of life [97]. The bone collagen was extracted following the Poznań Radiocarbon Laboratory protocol (see above) and subjected to isotopic determinations on bulk organic matter performed by the Stable Isotope Laboratory at Goethe University Frankfurt, Germany. Samples were processed using an element analyser (ThermoFisher, Flash 1112) connected to a continuous-flow inlet of a gas source mass spectrometer (ThermoFisher, MAT 253). Measurement quality is ensured by external analytical precision, usually better than ±0.2‰ for C isotopes and ±0.3‰ for N isotopes, and two-point corrections for both carbon and nitrogen isotopes by running USGS 24, IAEA-CH-7, IAEA-N1, and IAEA-N2 along with the samples.

### Food web and isoscape reconstruction

We use the grain samples for the cereal and the faunal samples for the animal source of the human food web in our analysis. The suitability of the *δ*^13^C and *δ*^15^N values of the three component groups – defined as cereals (subgroups *T. dicoccum*, *T. monococcum*); animals (subgroups herbivores, unknown); and humans – for modelling was tested by the Shapiro– Wilk test for normal distribution, using the software PAST, v. 4.11 [98]. Food webs were modelled in the software FRUITS, v. 3.1 [99,100] with the archaeological cereals and animals as food sources and the humans as target. We used the means of *δ*^13^C and *δ*^15^N of all charred grains (*T. dicoccum*, *T. monococcum*) and of the collagen of all animal bones and human skulls. The standard error of the mean (SEM) is considered as uncertainty. The relative proportions of food sources were calculated for the mean of individuals 5–7 as well as for the isolated bones. Offsets from diet to humans were set to 4.8‰ for *δ*^13^C and 5.5‰ for *δ*^15^N [99–103]. Uncertainties for individuals (0.3‰) and offsets (0.5‰) allow for uncertainties in the lab procedure and offset determination (S3 Tables).

For isoscape reconstruction, the mean isotopic signature of the plant diet was deduced by subtracting the offsets (4.8‰, 5.5‰) from the collagen values of the identified herbivore bones. We did the same for the omnivores and unidentified taxa, as these two subgroups of faunal remains do not exhibit any increased *δ*^15^N values that would point to substantial carnivory (S3 Table radiocarbon dating and stable isotopes). Reference ranges for *δ*^13^C in the plant component of the diet of herbivores in the forest-steppe and deciduous forests were calculated from published data adapted to differences in *δ*^13^C offsets [104] (5.1‰; 81) and atmospheric concentrations (7‰), with a *δ*^13^C concentration of 6.38‰ for Chalcolithic times taken from Ferrio et al. [105].

### Archaeological analogies

To help us interpret the significance of the Kosenivka human bone assemblage within the larger geographical framework of Chalcolithic communities of the forest-steppe area east of the Carpathians, we researched and re-evaluated finds of CTS human remains from settlement and burial contexts in the wider region, with a focus on the former, using both site-specific publications and compilations of human remains ([106–115,116], S5 Tables). We take into account analogies for burnt human remains from settlement contexts of the southeastern European Neolithic and Chalcolithic. By assessing the published number of human remains in relation to estimates of population sizes, we revive the discussion on the matter of the “missing dead”. We further evaluate the dating, find context, skeletal representativeness, and fragmentation of bone elements. The theoretical basis for the discussion on the formation processes of the mortuary deposits from Kosenivka are the observations of Weiss-Krejci [117].

## Results

### Human osteology

The collection of human bones recovered from the remains of house 6 and from its surrounding activity zone consists of ca. 50 commingled skeletal element fragments from at least seven individuals (for details and calculation of MNI, see S1 Table; for skeletal preservation, see S2 Appendix Fig 16). distributed across eight excavation squares (sq.), or 32 m^2^ (Fig 2). Except for one skull bone, all were located in the lower excavation layer, which revealed the inner structure of the house. For several bones, the exact find location is unknown. Only one bone, a patella, was preserved in its entirety. Some skeletal parts were found close to each other (e.g., those of individual 4), and a few fragments, some of which scattered within the feature structures, could be assigned to the same individual because they could be joined back together on fracture lines (individual 2, see Fig 2.C). The fact that there are cross-mends suggests post-mortem displacement and disturbance of the original, *in situ* situation. The skeletal element representation is broad; only hand, foot, and shoulder bones are missing (Fig 3; S2 Appendix Fig 17).

**Fig 2.**
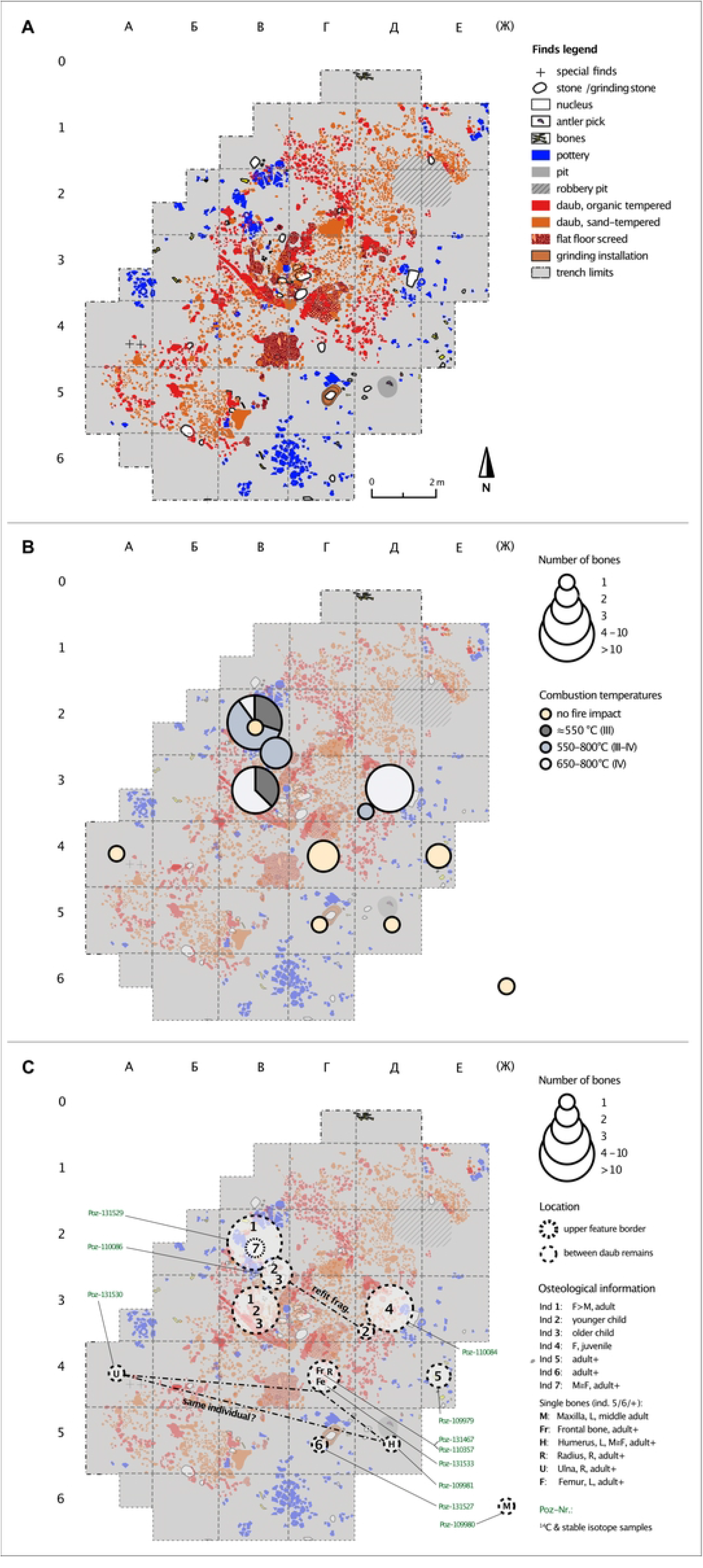
Kosenivka, excavation plans of house 6. Original plans modified after Kruts et al. [21]. (A) Archaeological finds. (B) Schematic localisation and number of human bones (centre of square) and degrees of fire impact, after Wahl [79]. (C) Contextual information, osteological results, and radiocarbon dating. Frag.=fragment, F=female, L=left, M=male, R=right. > means morphology tends to M respectively F, ≧ means morphology tends to M respectively F but sex identification is less reliable. For detailed information, see Tables 1 and 3, S1 Tables, and S2 Appendix Fig 17. Illustration: R. Hofmann, K. Fuchs.

**Fig 3.**
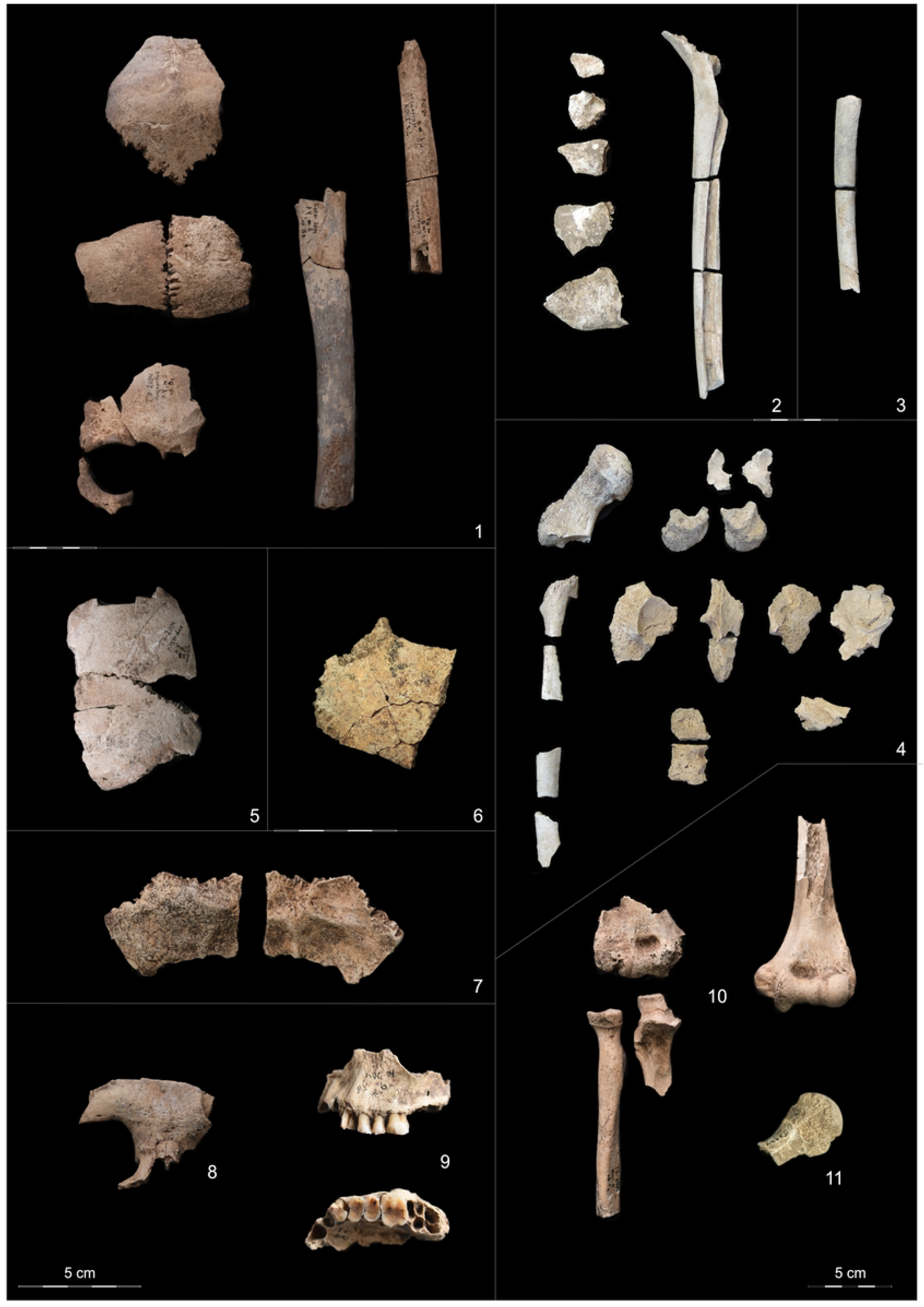
Kosenivka, selection of larger fragments from the human bone assemblage. (1–4) with fire impact. (1) Skull bones, femur and humerus diaphyses of individual 1, burnt. The beige colour is indicative of the burnt condition. (2) Skull fragments and femur diaphysis fragments of individual 2, calcined. (3) Femur diaphysis fragments of individual 3, calcined. (4) Femur, vertebra, pelvis, and sacrum fragments of individual 4, calcined. 5: Frontal and left parietal of individual 5. (6) Parietal fragment of individual 6. (7) Occipital (ecto- and endocranial views) fragment of individual 7. (8) Frontal bone fragment of individual 6+. (9) Left maxilla (buccal and palatinal views with teeth 23–26) of individual 5/6+. (10) Right and left (anterior views) arm bones from different locations but with matching anatomical features. Illustration/pictures: K. Fuchs, S. Storch.

**Table 1.** Summary of results from the osteological analyses per individual, current study and Kruts et al. [21]. Calcination grade (calc. grade) after Wahl [79]. For more detailed information, MNI rationale, and fire impact, see S1 Table and S2 Appendix Fig 17. Calc.=calcination; F=female; M=male; > means morphology tends towards M respectively F; ≧ means morphology tends to M respectively F but is less reliable; + means could represent an additional individual; ++ means unclear adult age; words in parentheses refer to thebone’s general appearance; n.d. means not determined. Square relates to the excavation unit as shown in the excavation plan (Fig 2). For other abbreviations, see Fig 2.

| Individual identifier | Individual identifier following Kruts et al. 2005 | Biological profile | Skeletal element(s) | Fire impact (calc. grade) | Square | Context |
| --- | --- | --- | --- | --- | --- | --- |
| 1 | 1 | F>M, <40 years, early adult | Skull and right humerus | Heavily burnt to calcined, III–IV | B2 | House centre |
|  | 1 | (early adult) | Right femur | Heavily burnt, III | B2–3 | House centre |
|  | 1 | >22 years (early adult) | Patella, tibia fragments, and fibula fragments | Heavily burnt to calcined, III–IV | B3 | House centre |
| 2 | 2 | Older child | Skull (different fragments) | Heavily burnt to calcined, III–IV | B2–3 | House centre |
|  | none |  | Rib fragments and femur fragment(s) | Heavily burnt to calcined, III–IV | B2–3 | House centre |
| 3 | none | Younger child | Left femur | Completely calcined | B2–3 | House centre |
| 4 | None (1, 3) | F≥M, 18–24 years, late juvenile | Pelvis, right femur, spine, and sacrum | Completely calcined | Д3 | Eastern part house centre |
| 5 | Male | n.d., 25–50 years, adult | Skull (left frontal and parietal) | No traces of fire | E4 | Eastern periphery |
| 6 | None | n.d.; 20–50 years, adult | Skull (left parietal) | No traces of fire | Г5 | Eastern house area |
| 7 | None | n.d., 20–50 years, adult | Skull (occipital) | No traces of fire | B2 | Western house centre |
| **Skeletal elements of unclear individual assignment** |  |  |  |  |  |  |
| 6, + | 3 | n.d., adult++ | Skull (frontal) | No traces of fire | Г4 | Eastern house area |
| 5, 6, + | None | n.d., 30–45+ years middle adult | Maxilla, left fragment, with teeth 24–26 | No traces of fire | (Ж6) | Overburden, no context |
| 5, 6, + | None | M≥F, adult++ | Humerus, left fragment | No traces of fire | Д5 | Eastern house area |
| 5, 6, + | None | n.d., adult++ | Radius, proximal fragment, right | No traces of fire | Г4 | Eastern house area |
| 5, 6, + | None | M≥F, adult++ | Ulna, right fragment | No traces of fire | A4 | Southwestern periphery |
| 5, 6, + | None | M≥F, adult++ | Humerus, right fragment | No traces of fire | No context | Unknown context |

The bones of individuals 1–4, represented by different skeletal parts (including skull, long bones, ribs, vertebrae, and pelvis), are heavily burnt (Table 1, Figs 3.1-4.B and 5). All remains exposed to fire were found within the concentrations of daub and other construction debris. The bones of individuals 5–7, represented by skulls and long bones only, are unburnt (Fig 3.5-11). The skeletal element representation suggests either heavy taphonomic damage to formerly articulated skeletons or intentional selection of body parts. These bones were recovered from the peripheral house activity area or the front part of the house feature itself. Burnt and unburnt bones were not found in close proximity to each other. The preservation varies from well preserved to fragile with respect to both firmness and surface condition. This is due to the different impact of fire as well as other taphonomic alterations.

**Fig 4.**
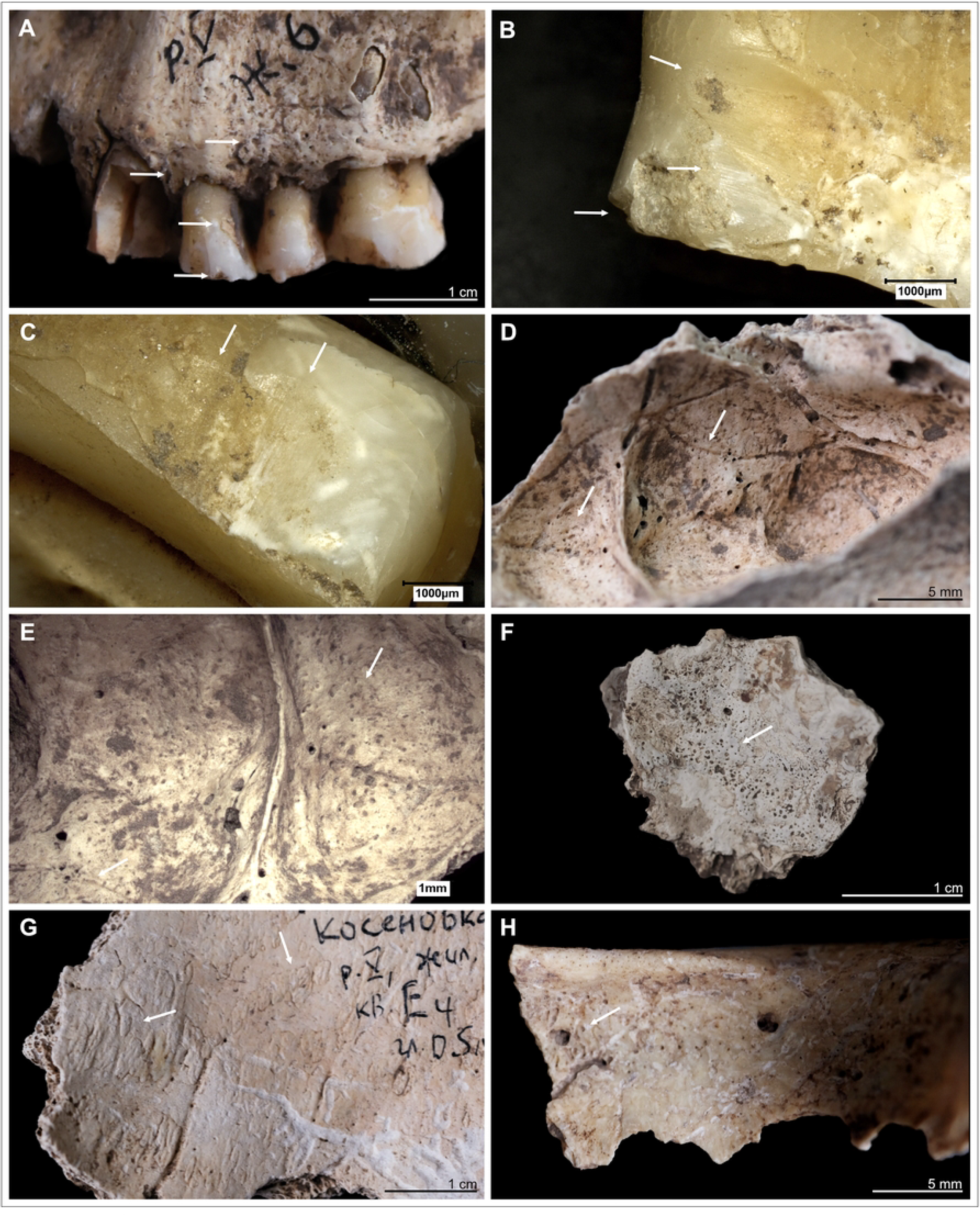
Kosenivka, selection of oral and pathological conditions. (A–E) Individual 5/6/+left maxilla. (A) Teeth positions 23–26 (buccal view). Signs of periodontal inflammation (upper arrows) and examples of dental calculus accumulation (third arrow) and dental chipping (lower arrow) on the first premolar (tooth 24). (B) First premolar (24, mesial view). Interproximal grooving with horizonal striations on the lingual surface of the root (upper arrow) and at the cemento–enamel junction (middle arrow). Larger chipping lesion (lower arrow). (C) Canine (23, distal view). Interproximal grooving, same location as on the neighbouring premolar (see B), but less distinct. (D, E) Signs of periosteal reaction on the left maxillary sinus (medio–superior view). Increased vessel impressions (D, upper arrow) and porosity, as well as uneven bone surface (D, lower arrow, E), indicating inflammatory processes. (F) Individual 2, left temporal, fragment (endocranial view). Periosteal reaction indicated by porous new bone formation (arrow). (G) Individual 5, frontal bone (endocranial view). Periosteal reaction indicated by tongue-like new bone formation and increased vessel impressions (arrows). (H) Individual 5/6/+, frontal bone, right part, orbital roof (inferior view). Signs of *cribra orbitalia* (evidenced by porosity, see arrow). Illustration: K. Fuchs.

**Fig 5.**
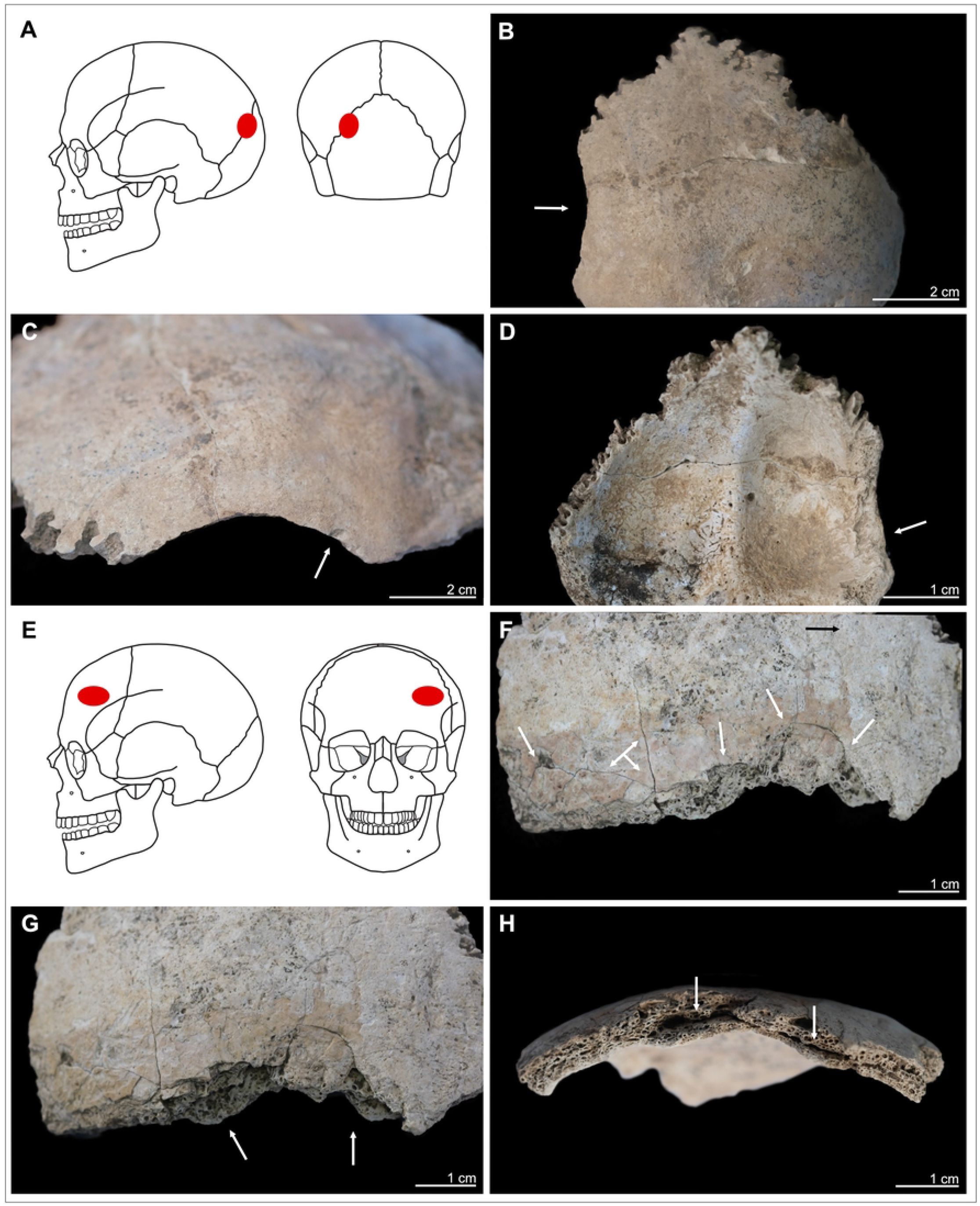
Kosenivka, selection of cases of perimortem cranial trauma, showing location and osteological details of the lesion. (A–D) Individual 1, occipital, left part. (A) Location of the trauma, on the left part of the back of the head. (B, C) *Lamina externa* (posterior and lateral views), showing an oval lesion (B, arrow) with a sharp rim and smaller punctual lesion (C, arrow). (D) *Lamina interna*, typical uneven, terraced appearance and enlarged rim of the lesion (arrow). E–H: Individual 5, frontal bone, left part, anterior view. (E) Location on the left forehead. (F) *Lamina externa* (anterior view), showing multiple fracture lines and terrace-like lesion rims (arrows). (G) *Lamina externa* (anterior–inferior view), showing the unevenly depressed rim of the lesion (arrows). (H) View from the diploe, showing the deformation of the cranial bone, with splitting inner cortex (arrows). Illustration: K. Fuchs.

Table 1 shows a summary of the MNI with their associated skeletal elements, isolated bones with possible individual association, impact of fire, and find context (for detailed information, S1 Table human remains and S2 Appendix human remains).

We estimate that least seven individuals are represented: two children of different ages (individuals 2 and 3); one late juvenile (18–20 years, probably female; individual 4); one middle-aged adult (ca. 25–40 years, probably female; individual 1); and three adults of undetermined age (likely of different ages but all between 20–50 years of age, two of them probably male; individuals 5–7). The remaining single bones most probably belong to two of the adult individuals (individuals 5 and 6) but may also represent additional individuals (+). The osteometric aspects of the left (sq. Д5) and right distal (unknown context) humerus epiphyses are almost the same (S1 Tables) and the proximal right ulna (sq. A4) and radius (sq. Г4) articulate very well with the right humerus. Thus, there is strong osteological indication that these four bones belong to the same individual, which would underline the vast dispersal of skeletal elements compared with a skeleton in anatomical articulation.

Although relatively few osteologic markers are available, the estimated biologic profile of the seven individuals appears to nearly correspond to a demographic cross-section.

Assuming that the assignments of the bones to different individuals are correct, it is noteworthy that the skeletal elements of the individuals found within the house are less scattered than of those of the individuals found in the periphery of the house. If secondary disturbance and taphonomy are responsible for this dispersal and also for the poor skeletal preservation indicated by skeletal element representation, this influence must have been stronger in the periphery than in the, perhaps more sheltered, surroundings of the burnt house structures. However, the locations of domestic finds, such as concentrations of sherds, and grindstones, and other tools, still allow us to discern activity areas.

### Palaeopathology: Disease and trauma

For a detailed summary of the descriptions that follow, as well as additional illustrations, see S1 Tables.

### Physical activity, inflammation, physical stress, and oral pathologies

The entheseal appearance of preserved ligament and tendon attachment sites does not indicate major biomechanical overloading of the musculoskeletal system for any of the individuals.

The same holds true for the joint surfaces. This could mean that (i) physical activities did not exceed physiological preconditions; (ii) the individuals were too young to have developed bone responses caused by repeated or permanent overexertion; or (iii) the body parts that showed overloading are not represented in the archaeological material.

The long bone remains of the upper and lower extremities, whether burnt or unburnt, do not show distinct responses to inflammation and other pathological changes. This finding may be influenced by the quite poor preservation of the bone surfaces. The cranial remains, however, do exhibit pathological changes, associated with periosteal reaction of the endocranium in the form of bone proliferation (individual 1, Fig 4.F) and increased vessel impressions, probably due to inflammatory processes (individual 5, Fig 4.G; S1 Tables). A mucous infection caused slight bone remodelling of the maxillary sinus of the adult individual (individuals 5, 6, or +; sinusitis, Fig 4.D-E; 93). The fragment also shows changes to the hard palate and the alveolar crests suggesting distinct inflammation of the stomal and periodontal tissues. The individuals (all adults) survived these processes long enough for the bone to have responded. A pathological indicator for unspecific physical stress was observed in the form of *cribra orbitalia* on the orbital roof of the single frontal bone fragment (sq. Г4; Fig 4.H). This distinct porosity, caused by an expansion of the haematopoietic tissue, is assumed to be a symptom for an increased production of red blood cells [119]. Red blood cells may be in high demand or biologically unavailable after someone has given birth or due to a number of different medical conditions, e.g., uterine diseases, long-term infections, scurvy, sickle-cell anaemia, or a deficiency in iron or other nutrients [118,120]. On this frontal bone, the *cribra orbitalia* has a rather smooth appearance and was probably an active process in childhood, adolescence, or early adulthood.

Macroscopic inspection of all four preserved teeth of the maxilla showed strong dental wear on the occlusal surfaces(grade 6 for premolars and grade 7 for molars, after Smith 1984; Fig 3.9, 4.A). The dentine portion is more eroded compared with the enamel portion, typically the result of intensively chewing foodstuffs with a highly abrasive consistency, such as cereals, and a rather acidic oral milieu (see [122]). Microscopic inspection identified faint traces of interproximal grooving on the interdental surfaces between the canine and the first premolar (Fig 4.B-C). Such grooves develop during repeated mechanical erosion of exposed tooth sections by penetration of a solid material, such as bone, wood, plant fibres, or sinew [123–125]. The grooves occur on the palatine tooth section inside the oral cavity and not on the buccal face. Another observed dental modification consists of small cracks in the enamel and dentine of the tooth crowns [126]. Considerable amounts of dental calculus had accumulated on all teeth, which reflects an alkaline pH level favourable for the mineralisation of dental plaque. Neither carious damage nor enamel developmental defects were observed.

### Cranial trauma

Two skull specimens show lesions typical for perimortem trauma. One traumatic injury was identified on the upper left section of the occipital bone of individual 1 (Fig 5.A-D). The roundish fracture line and the endocranial splitting of the bone are readily distinguishable from the other post-mortem damage to the occipital. The oval-shaped rim on the *lamina externa* is smooth and includes one very small area of depression on the lower part (3 × 2 mm) and adhering bone fragments. The lesion measures 30 × 6 mm, but as the corresponding bone parts are missing, the original dimensions remain unknown. The internal splitting of the *lamina interna* (Fig 5.D) indicates a high likelihood of the *dura mater* and brain tissue being injured, which would likely have been lethal during that time in the past. But since the pieces of crushed bone are not preserved, the clinical consequences remain unclear. An absence of signs of healing suggests that the individual survived this injury for a period of weeks at most [127], but probably died much earlier. The lesion was most probably caused by localised blunt force with a velocity high enough to split the three-layered cranial bone.

The forehead of individual 5 shows a fracture line on the left lateral section of the frontal bone that is atypical for post-mortem damage (Fig 5.E-H). The *lamina externa* is unevenly depressed, showing fine fracture lines of several mm in length and small bone parts dislocated towards the fracture line but still attached to the frontal bone, forming a terraced appearance. Although the *lamina externa* was clearly injured perimortem, the fracture line on the *lamina interna* has a split appearance suggestive of post-mortem damage. The diploe is clearly detached from the inner cortex, which is slightly deformed inwards (Fig 5.H). These characteristics are in line with typical signs for a depression fracture that did not open the cranial vault. In this type of cranial trauma, external forces impacting the skull were not strong enough to break through the elasticity of the spherically shaped cranial bone [58].

Since the remaining frontal bone is missing, the total size and severity of the lesion remains unclear. The depression is at least 38 mm in length along the fracture line and 18 mm in length towards the upper forehead (Fig 5.F-G). Again, there are no signs of healing. We cannot approximate the clinical implications of this injury; penetration of the *dura mater* is unlikely because the *lamina interna* is intact, but brain damage may have been caused by an endocranial haematoma. However, together with the lack of osseous responses indicating healing, this perimortem trauma had the potential to be lethal. Blunt forces are one main causation for such direct depression fractures.

Both traumatic injuries are localised above the head brim line, which is a decisive criterion for an interpersonal act of violence, rather than an accidental event or unintentionally executed violence [58,128]. For individual 1, the adult female, this means the strike probably happened while she and her attacker were face to face and was probably executed by a right-handed person. Individual 5 was hit on the back of the head. The injury has blurry borders in individual 5 and is quite large in individual 1. Spot-like blunt force induced by arrows is therefore unlikely. Trypillian stone tools, such as axes and daggers, may have been the weapons used. However, an accidental causation cannot be ruled out for either of these injuries. There is no evidence for traumatic injuries on any of the postcranial remains or the facial skeleton, which may be due to these skeletal elements’ relative underrepresentation in the human bone assemblage.

### Impact of fire and bioerosion

For a summary of diagnostic traits and their evaluation, see Table 2 and S1 Tables.

**Table 2.**
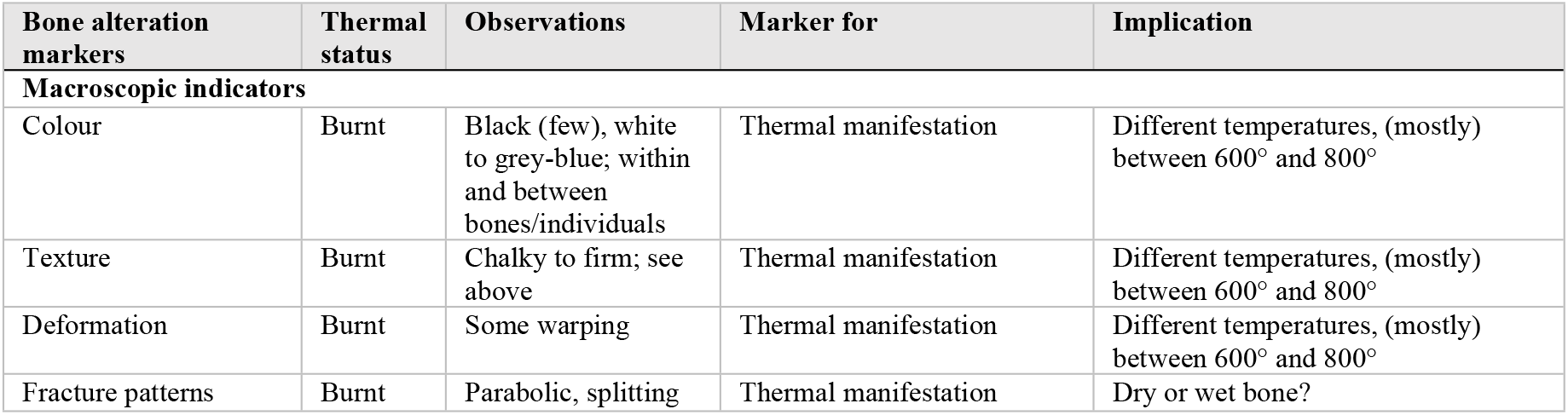
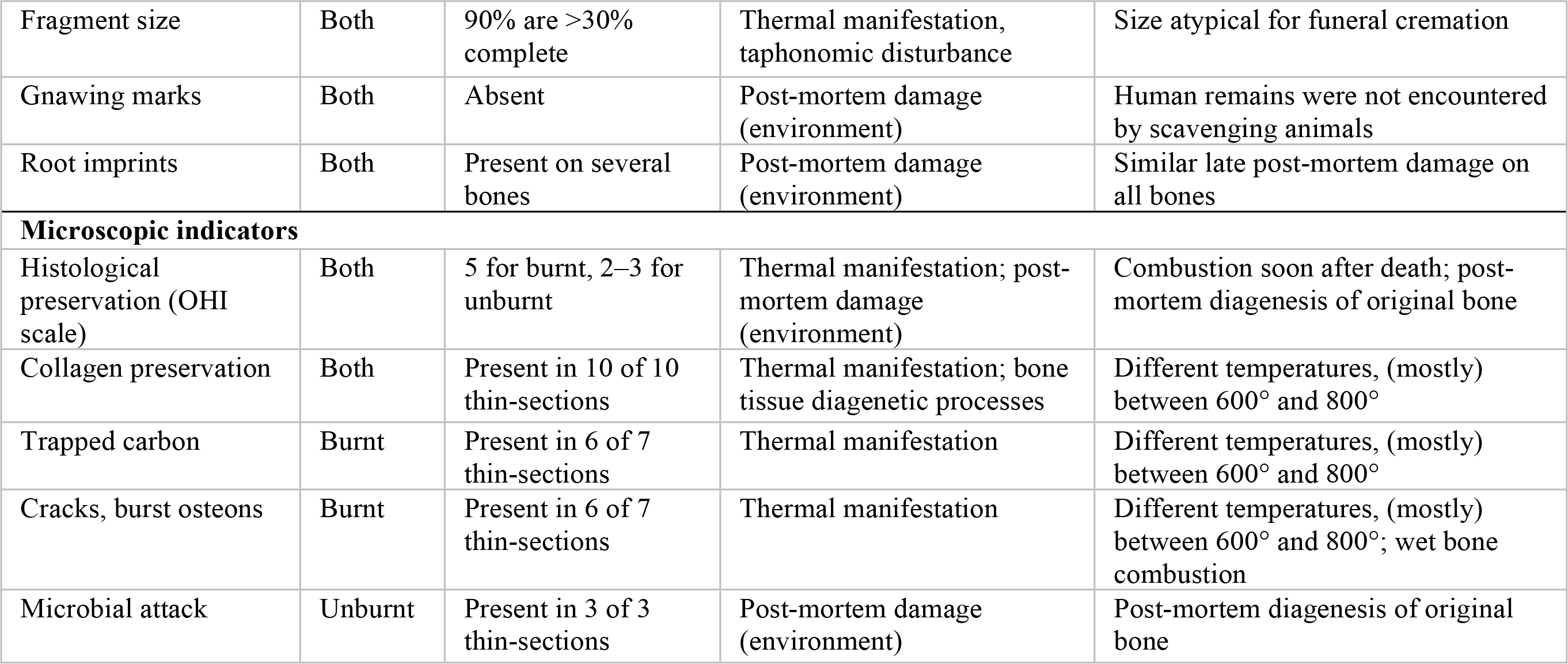
Summary of macroscopic and microscopic features of diagenetic bone alteration and their implications. For diagnostic marker descriptions and implications, see Ellingham et al. (78); Wahl (79). For more examples, detailed descriptions, and illustrations, see Fig 6 and S1 Tables.

| Bone alteration markers | Thermal status | Observations | Marker for | Implication |
| --- | --- | --- | --- | --- |
| **Macroscopic indicators** |  |  |  |  |
| Colour | Burnt | Black (few), white to grey-blue; within and between bones/individuals | Thermal manifestation | Different temperatures, (mostly) between 600° and 800° |
| Texture | Burnt | Chalky to firm; see above | Thermal manifestation | Different temperatures, (mostly) between 600° and 800° |
| Deformation | Burnt | Some warping | Thermal manifestation | Different temperatures, (mostly) between 600° and 800° |
| Fracture patterns | Burnt | Parabolic, splitting | Thermal manifestation | Dry or wet bone? |
| Fragment size | Both | 90% are >30% complete | Thermal manifestation, taphonomic disturbance | Size atypical for funeral cremation |
| Gnawing marks | Both | Absent | Post-mortem damage (environment) | Human remains were not encountered by scavenging animals |
| Root imprints | Both | Present on several bones | Post-mortem damage (environment) | Similar late post-mortem damage on all bones |
| **Microscopic indicators** |  |  |  |  |
| Histological preservation (OHI scale) | Both | 5 for burnt, 2–3 for unburnt | Thermal manifestation; post-mortem damage (environment) | Combustion soon after death; post-mortem diagenesis of original bone |
| Collagen preservation | Both | Present in 10 of 10 thin-sections | Thermal manifestation; bone tissue diagenetic processes | Different temperatures, (mostly) between 600° and 800° |
| Trapped carbon | Burnt | Present in 6 of 7 thin-sections | Thermal manifestation | Different temperatures, (mostly) between 600° and 800° |
| Cracks, burst osteons | Burnt | Present in 6 of 7 thin-sections | Thermal manifestation | Different temperatures, (mostly) between 600° and 800°; wet bone combustion |
| Microbial attack | Unburnt | Present in 3 of 3 thin-sections | Post-mortem damage (environment) | Post-mortem diagenesis of original bone |

**Table 3.**
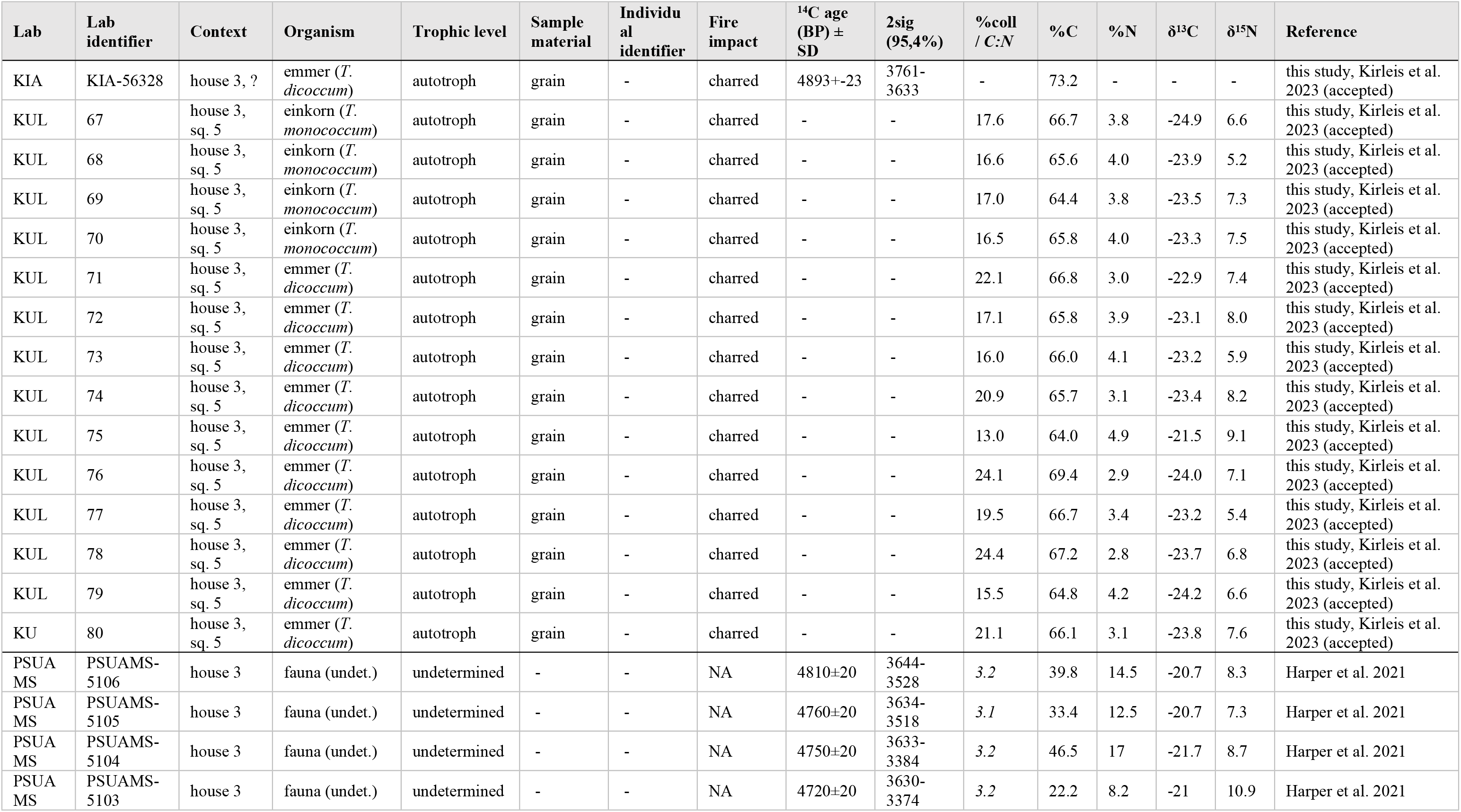
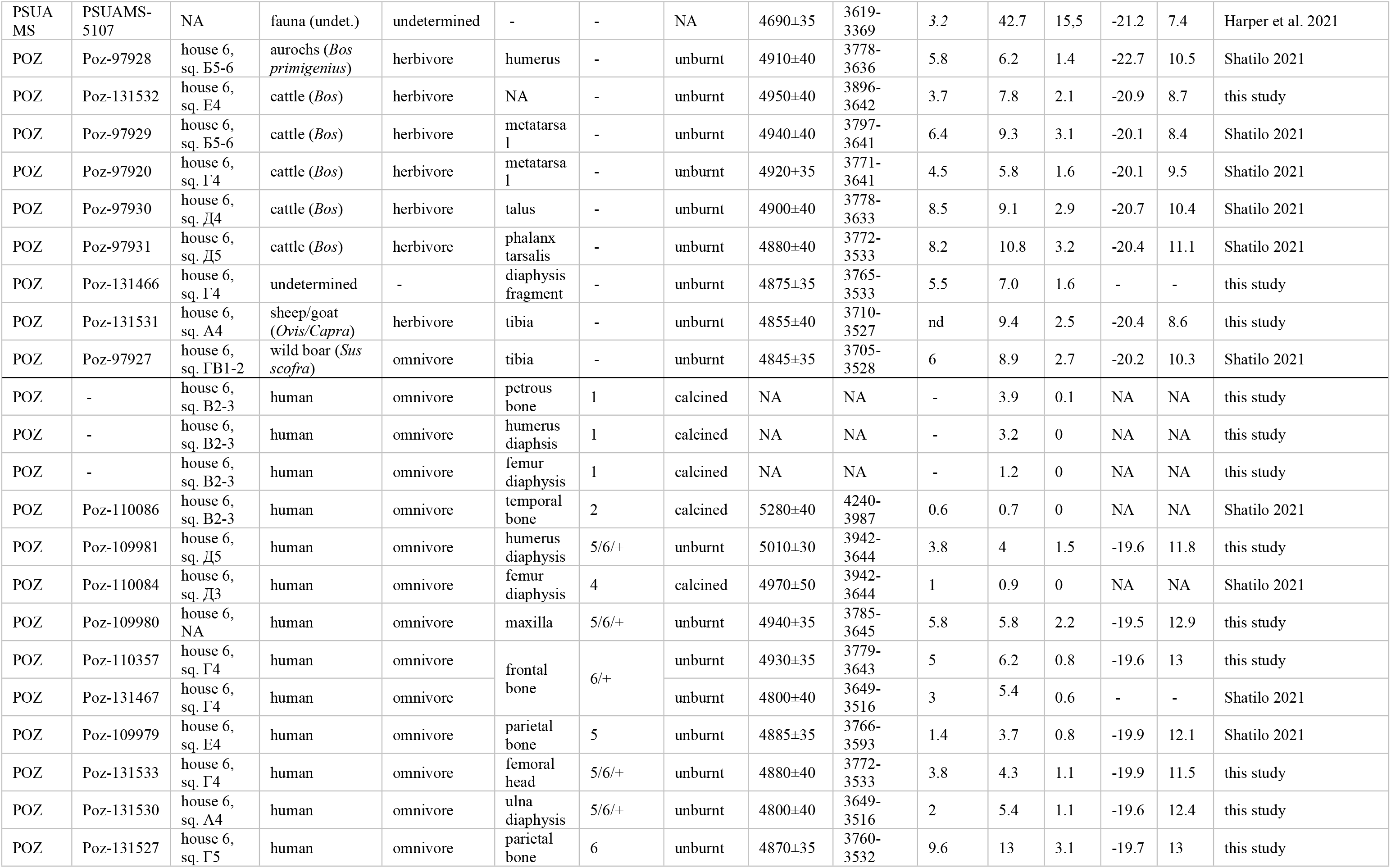
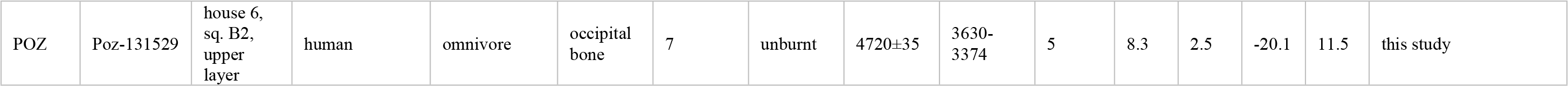
Sample characteristics and results for the radiocarbon and carbon and nitrogen stable isotopic analyses of plant, animal, and human remains, including published data (S3 Tables). KIA=Leibniz Labor für Altersbestimmung und Isotopenforschung / Leibniz Laboratory for Radiometric Dating and Stable Isotope Research, Kiel University, Germany; PSUAMS=Radiocarbon Laboratory, Pennsylvania State University, USA; KUL=Katholieke Universiteit Leuven (Catholic University of Louvain), Belgium; POZ=Poznańskie Laboratorium Radiowęglowe / Poznań Radiocarbon Laboratory, Poland. Ind.=individual; sq.=square.

### Impact of fire

All of the bones that exhibit considerable evidence for the impact of fire were found in the house centre (squares B2–B3, Д3). Macroscopic and microscopic markers of thermal manifestations point to exposure to temperatures of 550–800°C (Figs 6-7, Table 2; [64]). Bone colour, texture, and deformation vary within the same specimen and between individuals (Fig 6, S1 Tables) indicating that the degree of temperature or exposure time were inconsistent. The majority of the specimens show evidence for combustion at temperatures that exceeded 650°C (Table 1-2, S1 Tables). It should be noted that some researchers assign higher temperatures for the observed changes (between 800°C and 1000°C; [53]).

**Fig 6.**
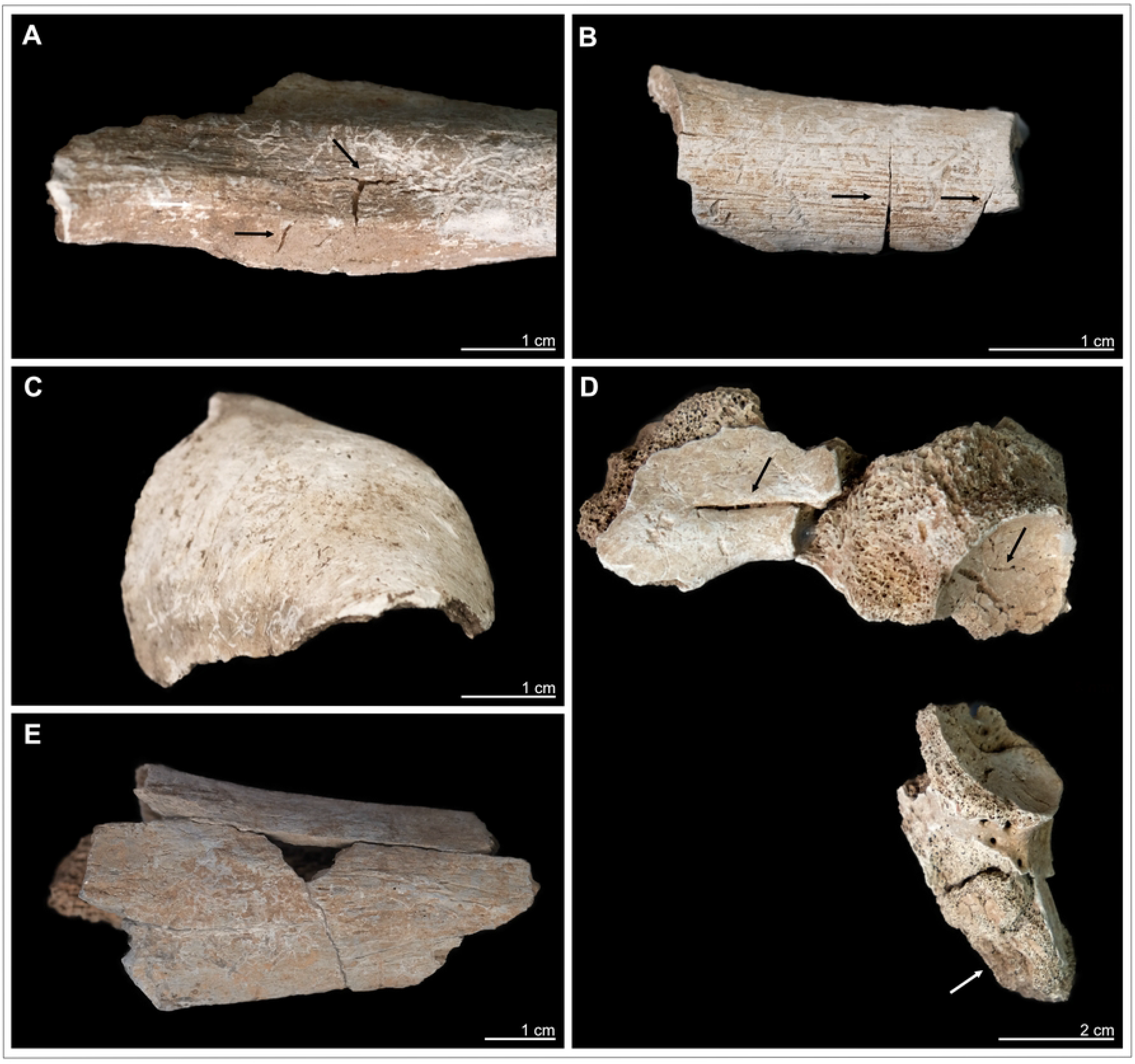
Kosenivka, examples of macroscopic features of thermal impact on larger human bone fragments that are atypical for cremation as a funeral practice. (A, B, D) Fracture patterns, splitting and heat cracks (arrows). (C, D) Distinct deformation (warping) as typically occurs under higher combustion temperatures (left arrow). (A–E) Discoloration to white and to grey-blue colour, demonstrating different combustion temperatures. (A) Individual 3, left femur diaphysis, proximal (medial view). (B) Individual 2, left femur diaphysis, distal (lateral view). (C) Individual 2, right parietal bone (anterior view). (D) Individual 4, right iliac and ischiatic bones (lateral view; arrow: unfused apophysis ischiatic surface). (E) Individual 1, right tibia diaphysis, proximal (posterior view). For more information, see S1 Tables and S2 Appendix. Illustration: K. Fuchs.

**Fig 7.**
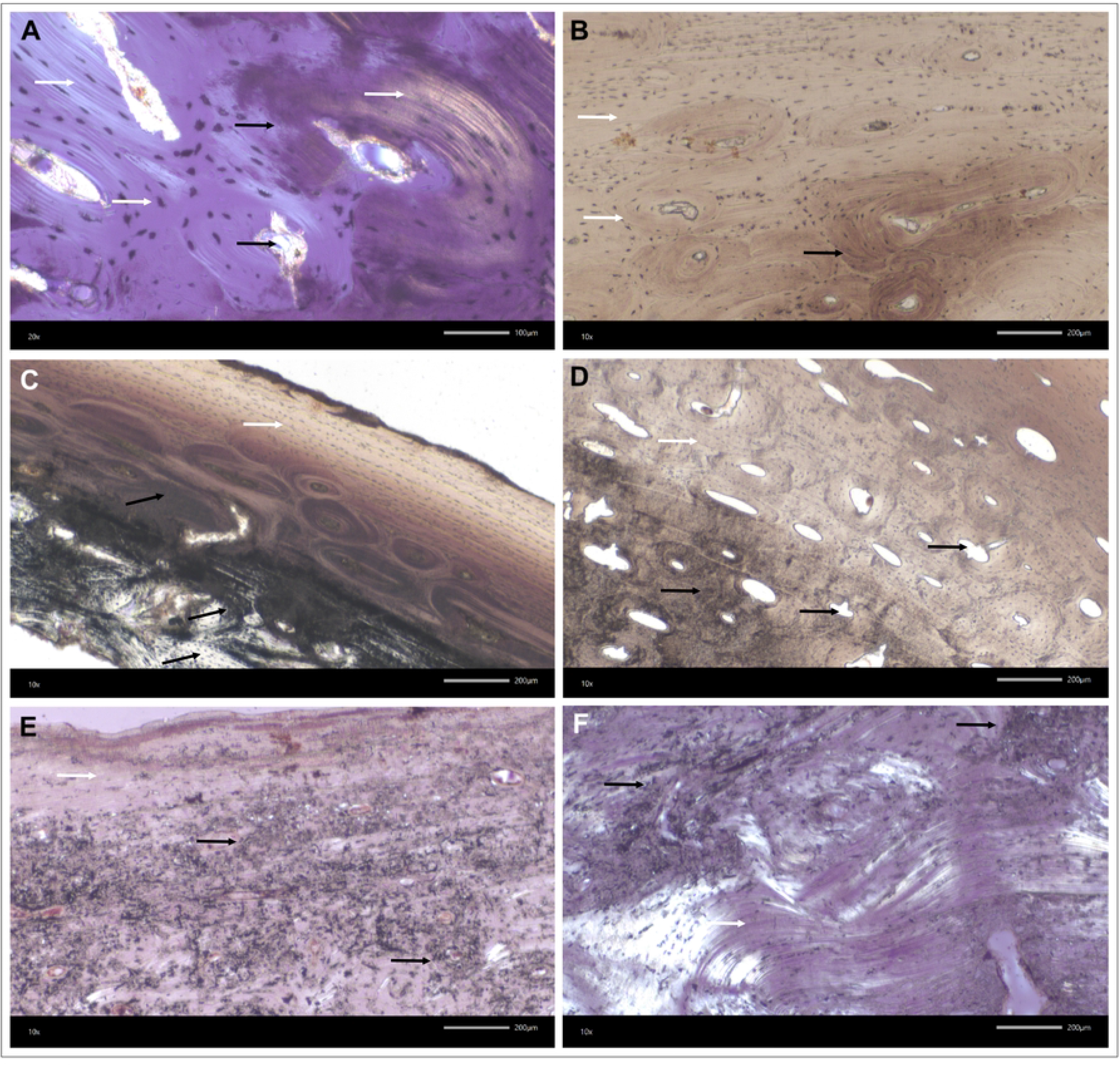
Kosenivka, transmitted light microscopy of bone thin-sections. Microscopic features of preservation, fire impact and bioerosion in bone microstructure. (A–D) Calcined bone. Black arrows indicate thermal impact, that is, calcined bone with discoloration by trapped carbon, burst haversian canals, and reduced osteocyte distance due to shrinking. White arrows indicate well-preserved microstructures, such as osteons, osteocyte lacunae, circumferential lamellae, and collagen (A, yellow colour). (A) Individual 1 (adult), right femur, proximal diaphysis, OHI of 5 (transversal section, polarised light). (B) Individual 1, left parietal, cortical bone of lamina interna, OHI of 5 (vertical section, plain light). (C) Individual 2 (younger child), left parietal, OHI of 5 (vertical section, plain light). (D) Individual 3 (older child), left femur, diaphysis, OHI of 5 (transversal section, plain transmitted light). (E, F) Unburnt bone with poorly preserved bone histomorphology. Strong impact of microbial attack visible by focal deconstruction (e.g., longitudinal tunnelling; black arrows; see S2 Appendix Fig 17.5-7) and a few well-preserved, original areas (white arrows). (E, F) Individual 5/6/+ (adult), right humerus, distal epiphysis, OHI of 3 (transversal sections, polarised light). For more examples and age-related histomorphological features, see S2 Appendix Fig 17.

Morphological features, such as warping, splitting cracks, or parabolic fractures, were observed on several fragments (Fig 6.A-B.D). These features are characteristic for fresh bone that has been burnt but can also appear on dry bone exposed to high temperatures [129]. The extent of heat-induced shrinkage cannot be determined, yet small spaces between osteocyte lacunae show considerable bone shrinkage due to thermal impact. This makes it difficult to evaluate osteological markers for biological age and age-at-death. Most of the skeletal elements are more than 30% complete. Small and very small fragments are relatively rare. This level of fragmentation is often observed in unfavourable preservation conditions and is atypical for intentional cremation as a funeral practice [72], a practice that is well known for the Trypillia [130].

Overall, histological preservation of the analysed burnt bones is very good (Fig 7.A-D; OHI of 5, S1 Tables, S2 Appendix Fig 17), showing the authentic matrix, with well-identifiable microscopic units for cortical and cancellous bone tissue, such as the cylindrical osteons (cortical bone), osteocytes (partly with *canaliculi*), and circumferential lamellae. This preservation gives us additional information for estimating the age-at-death of the individuals these specimens derive from (S1 Tables). Furthermore, intra-bone differences in processes of combustion and temperature impacts are well observable through carbon discoloration in the inner compacta and trabecular structures (indicating combustion of organic compounds), partial preservation of collagen structures, and completely calcined regions (Fig 7.A). Markers for fresh bone combustion, such as burst Haversian canals and heat cracks, were identified in five of the seven histological thin-sections from burnt specimens.

### Histotaphonomy and bioerosion

The histological preservation of the unburnt bone tissue is poor (OHI 2–3) compared with that of the burnt bone (OHI 5; see Fig 7.A-D). Microbial focal deconstruction and carbon inclusion, even in osteocytes, are clearly distinguishable. Signs of microbial attack in the form of Wedl, linear longitudinal, and budded tunnelling were observed on all tissue thin-sections of unburnt specimens and in different skeletal elements (Fig 7.E-F, S2 Appendix Fig 17.5-7; see [44,45]). The extent of microscopic damage is relatively great, which means the collagen-rich bone had sufficient exposure to the causative microbial agents, in terms of both time and environmental conditions. Markers of microbial attack were absent in the burnt specimens, indicating that the time between the death of these individuals and the combustion of their cadavers was not sufficient for microbes to have migrated into the bone matrix and fed on its organic matter.

Estimating the onset of this natural diagenetic path of putrefaction is complex, and results from experimental studies are largely not transferable to an archaeological situation like that at Kosenivka [64,131]. There is consensus that the post-mortem environment (humidity, temperature, oxygen availability, microbial abundance and diversity) and the condition of the cadaver itself (whether intact, injured, defleshed) have strong impact on bio-erosive processes (for a summary, see [103]). Estimates of the amount of time that has to elapse after death for microorganism activity to be visible in the bone might vary from days to several years; this also depends on whether enteric (during putrefaction) or exogenous (soil organisms) osteolytic microbiota attack the skeletal body [62]. Lemmers and colleagues [129] emphasise that an absence of bioerosion is not evidence of combustion directly following death, but that its presence is evidence for a subsequent treatment including fire. Yet, the extensive microscopic bone damage observed in the unburnt specimens from Kosenivka suggests that the depositional environment of house debris, remains of domestic artefacts, and natural soils was favourable for microbial activity. It remains unclear whether this activity took place before or (long) after the burning of the house.

### Further taphonomic indicators

Both burnt and unburnt bone surfaces exhibit later post-mortem erosive damage due to mechanical impact and root penetration. Gnawing marks are completely missing, meaning that the human remains were not encountered and displaced by scavenging animals.

### Radiocarbon dating

Sample characteristics and radiocarbon results for the human and animal bone are summarised in Table 3 (see S3 Tables and S4 Appendix).

The 26 available ^14^C dates were measured in three different laboratories, the results of one of which differ slightly in average age, but not considerably. Overall, the uncalibrated ^14^C age spans from 5280±40 (Poz-11086, skull bone individual 7, house 6) to 4690±35 (PSUAMS-97972, faunal bone from an undetermined domestic feature; Table 3). The sum calibration (2σ) of dates obtained for house 6 measured by the Poznań laboratory spans a maximum range of 4240–3374 cal BCE. The date from the Leibniz laboratory for the cereal grain from house 3 falls within this range as well (KIA-56328, 3710–3636 BC, 2σ, Fig 8). However, the sum calibration of the other five samples from house 3 results in a slightly younger age, with a maximum span of 3644–3374 cal BC (2σ, Fig 8). Therefore, we attribute these results to inter-laboratory discrepancy.

**Fig 8:**
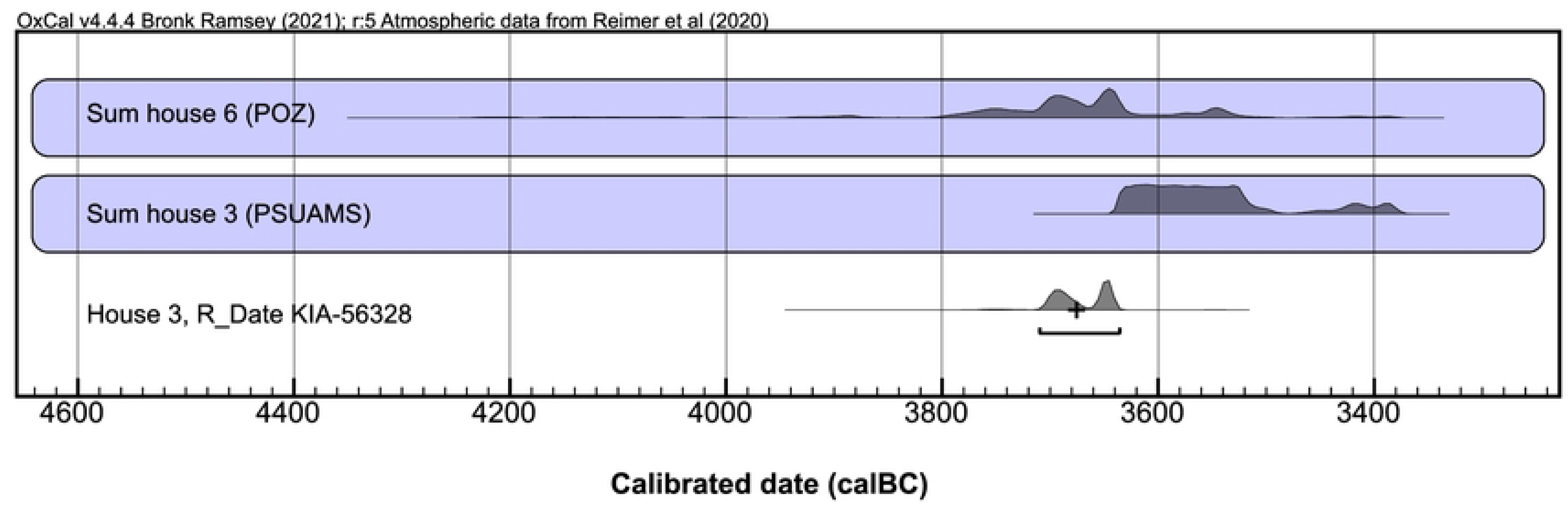
Multiplot of sum and single calibrations of samples per laboratory. House 6, human and faunal bone samples (n=20), POZ=Poznań Radiocarbon Laboratory, Poland. House 3, faunal bone samples (n=4), PSUAMS=Pennsylvania State University AMS Laboratory, USA. House 3, charred emmer grain (n=1), KIA=Leibniz Laboratory for Radiometric Dating and Stable Isotope Research, Kiel University, Germany. Illustration: K. Fuchs.

### House 6

The 20 bone samples comprise 11 human specimens (2 of them burnt) from at least 5 individuals, and 9 faunal specimens (Table 3). The two burnt samples only just passed the collagen quality control criterion, with 0.7% and 0.9% yield, but since AMS dating of calcined material is based on the structural carbonate, the results can be considered valid (Poz-110084, Poz-110086; 65). All unburnt bone samples passed the quality control, and the results do not indicate a correlation between collagen yield and ^14^C age (see S4 Appendix, Fig 18). Based on the carbon and nitrogen isotopic values, we can exclude a major reservoir effect resulting from the consumption of aquatic food sources (see next section).

After calibration, the human and faunal sample groups do not differ statistically; thus a comparable dating can be assumed (Fig 9). One calcined sample (individual 2, Poz-11086) pre-dates the other human samples. Due to the absence of stratigraphic evidence for this bone representing an older deposition, we argue that there is likely an offset in this date, explained by the ‘old-wood effect’ [81,132], in this case presumably the wooden parts of the house construction [11]. In contrast, the oldest date on the occipital bone from individual 7 (Poz-131529, 3630–3374 cal BCE, 2σ) coincides with its stratigraphically younger age, it having been found in an upper layer. Thus, this bone could represent a later deposition. Apart from these two dates from individuals 2 and 7, the dates from the other samples, including the faunal specimens, resulted in a statistically consistent unmodelled time range of ca. 3800– 3600 cal BC. Further interpretation is influenced by wide-spanning probabilities and wiggles in the calibration curve (S4 Appendix Figs 19-20), as underlined by a high intra-individual variability of the mentioned double sampling of the frontal bone, which generated results of 3779–3643 and 3649–3516 cal BC (Poz-110357, Poz-131467, 2σ). The same argument, relating to wide-spanning probabilities and wiggles, applies to the two arm bone samples (a left humerus and a right ulna), as their anatomical markers indicate that they belong to the same individual (Poz-109981, Poz-131530; see S1 Tables).

**Fig 9.**
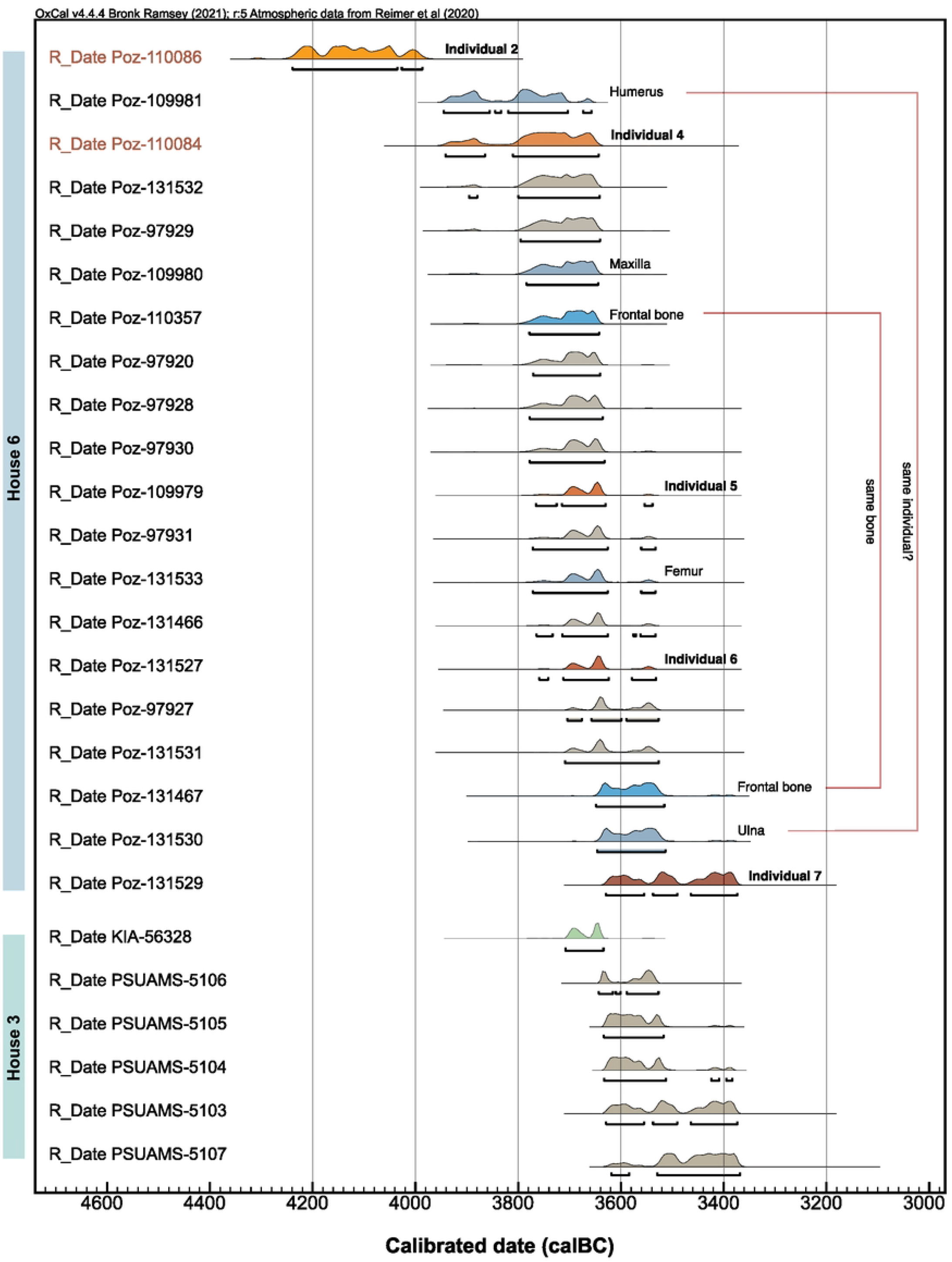
Multiplot of the radiocarbon dates of human, faunal, and botanical (cereal) samples from houses 3 and 6 at Kosenivka. Obtained by using OxCal, v. 4.4.165 (74; in conjunction with the INTCAL20 calibration curve). Probability range is 95.4% (2σ). Red sample identifier font indicates calcined bone sample. Light brown–coloured probability distributions mark the faunal samples, green the cereal sample, orange to dark red the identified human individuals, and blue the isolated human bone elements (see Table 3). Illustration: K. Fuchs.

To improve the interpretative potential of the results, we applied Bayesian modelling (Fig 10; S4 Appendix Fig 20), considering the one bit of stratigraphic information and treating the human and faunal samples as two separate assemblages (see Materials and methods).

**Fig 10.**
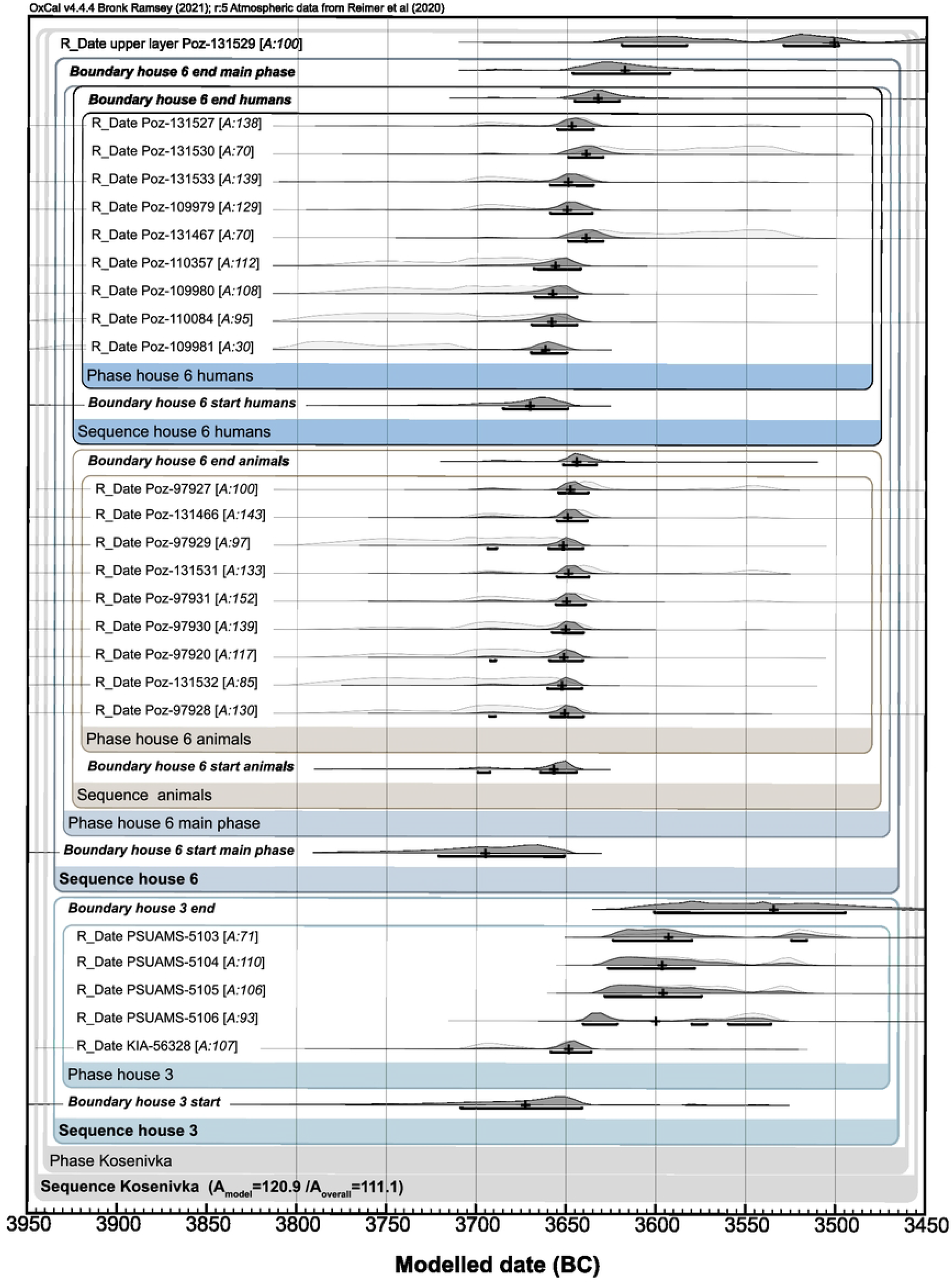
Bayesian modelling of the radiocarbon dates. We used the OxCal v. 4.4.165 functions “sequence,” “boundary,” and “combine” for considering archaeological context and stratigraphic information (74; in conjunction with the INTCAL20 calibration curve). The probability range (dark grey) is 68.3% (1σ). The individual agreement with the overall model is indicated by the A value (%) (in square brackets) per sample (should be >60%). For the modelling code, see S3 Tables; for the curve plot, see S4 Appendix Fig 20. Illustration: J. Müller, K. Fuchs.

**Fig 11.**
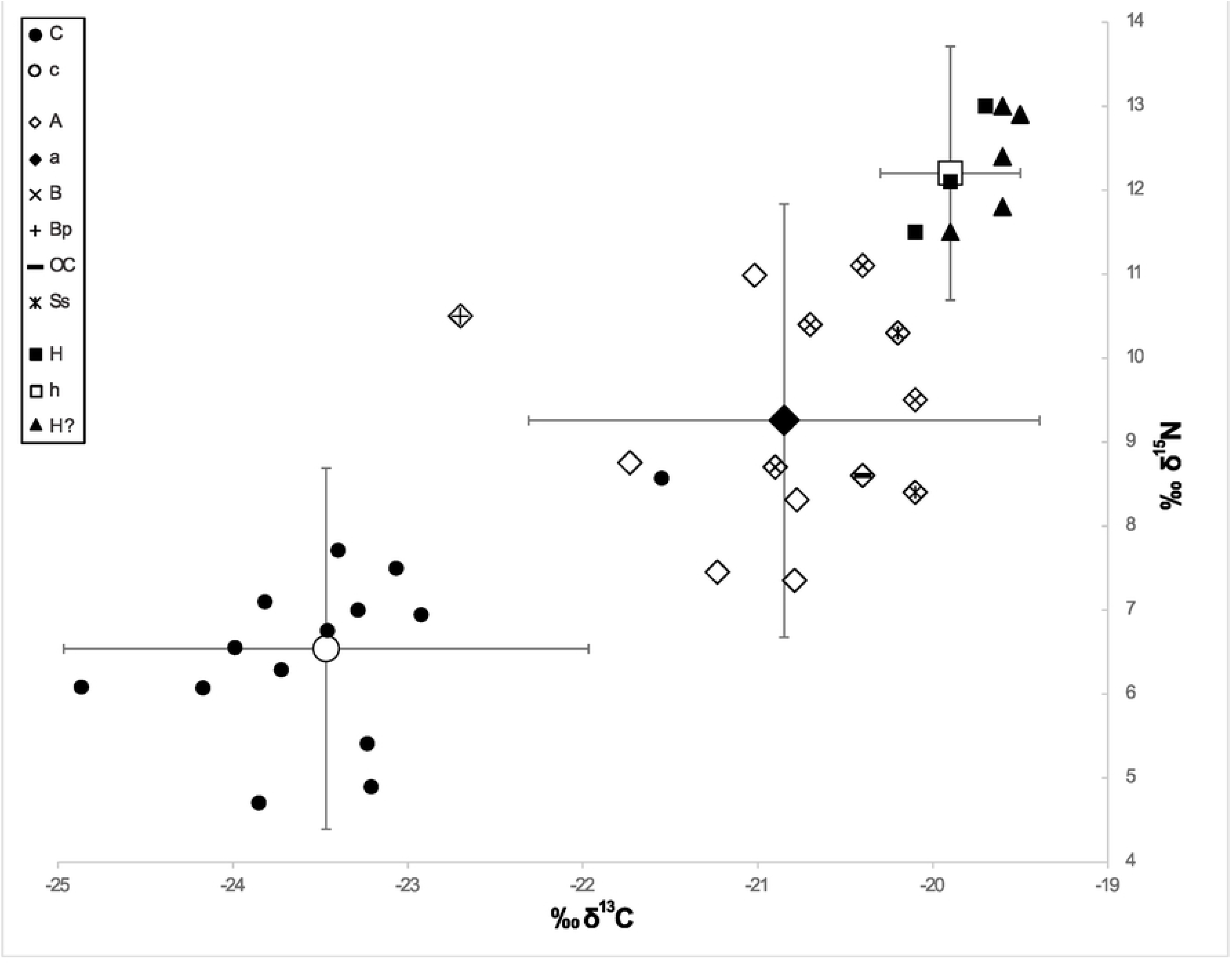
δ^13^C and δ^15^N values from Kosenivka. Unburnt cranial remains representing individuals 5–7 (H), animal bones (A), and cereal grains (C) and their respective means (h, a, c), with error bars representing 2 SD. Human bones with unclear individual affiliation are plotted as triangles (H2). δ^15^N of cereals has been reduced by 0.5‰ for charring. B=*Bos taurus*; Bp=*Bos primigenius*; OC=*Ovis aries/Capra hircus*; Ss=*Sus scrofa*. Taxonomic identity of the remaining animal bones not determined. Illustration: F. Schlütz, K. Fuchs.

Sequences were created by separating (i) the youngest date, with the stratigraphically younger age; (ii) the human samples with the wider age ranges; and (iii) the faunal samples (OxCal functions “boundary,” “phase,” and “sequence,” calculating 1σ and median values; Fig 10). The model generated a high statistical significance (^A^model/^A^overall 120.9/111.1). Poz-109981, one of the arm bones, does not fit this model with A<30%. But since the two dates from the same frontal bone failed when applying the OxCal function “combine,” we consider these deviations to be the result of analytical bias. The main results are:

- The animals and humans (individuals 3, 5, and 6, ++) represented by the analysed samples died ca. 3690–3620 cal BC (calculated median duration 79 years).
- The animals died ca. 3660–3645 cal BC (calculated median duration 11 years), the humans (individuals 3, 5, and 6, ++) ca. 3670-3635 cal BC (calculated median duration 37 years).
- The human skull bone, from individual 7, was probably deposited on the upper layer of the already burnt house; thus individual 7 died ca. 3620–3380 cal BC (median 3500 cal BC).

The shorter intervals for the animals can be explained by their shorter lifespans; cattle, for instance, were usually slaughtered when 10 years old, whereas eight out of nine samples in this modelling stem from adult humans, whose lifespan would have been considerably greater than that.

Overall, the data are statistically close enough that the calculated lifetimes of the animal and human individuals associated with house 6 are probably more consistent with the human results than with the combined human and animal results. It is very likely that individuals 4 to 6, plus potentially those represented by isolated bones (5/6++), died during the house’s occupation. Individual 7 died ca. 130 years after the other individuals, and its skull bone was probably deposited after the house had already been abandoned. We can thus conclude that house 6 was in use ca. 3660–3635 cal BC (1σ), a timespan that includes the median duration of the lifespan of the animals and of the most recent of the humans.

### House 3

As mentioned above, the date of the cereal grain analysed at the Leibniz laboratory fits with the dates obtained for house 6, while the dates of the faunal bones analysed at the Penn State laboratory have a longer probability span, towards a younger age. Altogether, the samples for house 3 date to ca. 3675–3535 cal BC. If we give more significance to the cereal grain date, since there is a strong indication of laboratory bias for the other samples, we can conclude that house 3 was in use ca. 3702–3640 (median 3680) cal BC and therefore contemporary with house 6 (Fig 10).

#### The Settlement of Kosenivka

On the basis of the dates measured at the Leibniz and Poznań laboratories, we conclude that houses 3 and 6, including the humans, animals, and cereals associated with them, existed approximately contemporaneously, between 3700 and 3620 BCE. As we lack prior information, especially with respect to stratigraphy and ritual activities, it is not possible to narrow the time span to less than 120 years. Further activities took place at the site several generations later (the deposition of the skull bone and of the faunal bone from an unknown context; [8]), probably around 3500 cal BC. Thus, our results place the settlement of Kosenivka in Late Trypillia C1 and some activities at the site in C2, but mainly in the final phase of Middle Trypillia. Overall, these results support the hypothesis of Kruts and colleagues [21] that the humans represented by the skeletal remains found in house 6 died together, in a single event, such as a fire event.

### Stable C and N isotopes: Flora, fauna, and humans

#### Stable C and N isotopes from cereals and bones

The distribution of *δ*^13^C values for the sample categories cereals, human bones, and undetermined animals is not significantly different from normal (Shapiro–Wilk test p>0.05; Table 4). The *δ*^13^C values of the animals as a whole and of the subgroup herbivores exhibit a significantly non-normal (p<0.05) distribution, but if we remove the extreme *δ*^13^C value of *B. primigenius*, they exhibit a normal distribution (animals p=0.505, herbivores p=0.425). The *δ*^15^N values exhibit a normal distribution within each group and subgroup.

**Table 4.** Summary of the means, standard deviations (SD), ranges, and results of Shapiro–Wilk test of δ^13^C and δ^15^N for the groups cereals, animals, and humans and their respective subgroups. For the categories animals and herbivores, a non-normal distribution of the δ^13^C values (p<0.05) is indicated by a *. Standard error of the mean (SEM) for groups is that used in the FRUITS food web model. ** δ^15^N of cereals adjusted by −0.5‰ for charring.

| | | $\delta^{13}\text{C}$ | | | | | $\delta^{15}\text{N}^{**}$ | | | | |
| --- | --- | --- | --- | --- | --- | --- | --- | --- | --- | --- | --- |
| | | Mean ± SD | SEM | Range | Shapiro–Wilk test | | Mean ± SD | SEM | Range | Shapiro Wilk test | |
|  | N | [‰] | [‰] | [‰] | W | p | [‰] | [‰] | [‰] | W | p |
| **Cereals** | 14 | -23.47 ± 1.50 | ± 0.201 | -24.87 to -21.55 | 0.92 | 0.234 | 6.54 ± 2.15 | ± 0.287 | 4.70 to 8.57 | 0.98 | 0.963 |
| *T. dicococcum* | 10 | -23.31 ± 1.48 | - | -24.17 to -21.55 | 0.88 | 0.118 | 6.70 ± 2.20 | - | 4.90 to 8.57 | 0.99 | 0.999 |
| *T. monococcum* | 4 | -23.87 ± 1.41 | - | -24.87 to -23.29 | 0.88 | 0.349 | 6.13 ± 2.06 | - | 4.70 to 7.00 | 0.90 | 0.414 |
| **Animals** | 13 | -20.85 ± 1.46 | ± 0.202 | -22.70 to -20.10 | 0.87 | 0.049* | 9.26 ± 2.58 | ± 0.357 | 7.35 to 11.10 | 0.92 | 0.267 |
| Herbivores | 7 | -20.76 ± 1.81 | - | -22.70 to -20.10 | 0.74 | 0.009* | 9.60 ± 2.15 | - | 8.40 to 11.10 | 0.90 | 0.311 |
| *Sus scrofa* | 1 | - | - | -20.2 | - | - | - | - | 10.3 | - | - |
| undetermined | 5 | -21.11 ± 0.79 | - | -21.73 to -20.78 | 0.88 | 0.316 | 8.57 ± 2.94 | - | 7.35 to 10.99 | 0.86 | 0.225 |
| **Humans** | 8 | -19.90 ± 0.40 | ± 0.115 | -20.10 to -19.70 | 0.89 | 0.242 | 12.20 ± 1.51 | ± 0.436 | 11.50 to 13.00 | 0.99 | 0.780 |

The isotopic values of the cereals range from −24.9‰ to −21.5‰ (mean −23.5‰) for *δ*^13^C and from 4.7‰ to 8.6‰ (6.5‰) for *δ*^15^N. Those for animals range from −22.7‰ to −20.1‰ (−20.9‰) and from 7.4‰ to 11.1‰ (9.3‰). Those for human bone collagen range from −20.1‰ to −19.7‰ (mean −19.9‰) for *δ*^13^C and from 7.4‰ to 11.0‰ (mean 8.6‰) for *δ*^15^N (Table 3, Fig 11). The *δ*^13^C values of the cereal grains therefore fall within the normal range of C_3_ plants [133], but their *δ*^15^N seems to be relatively high, pointing to a supply of nitrogen that exceeds what would have been naturally available in the soil [89]. The range in *δ*^13^C values of the animal bone collagen closely resembles that of the mega-sites of Maidanetske and Nebelivka, and represents open pastures to woodlands [13,36,134]. The high *δ*^15^N values also resemble the values from Maidanetske and Nebelivka, indicating intensive pasture practices. Such high values were not seen in the small-sized Stolniceni data set [13].

For δ^15^N, the range, between 11.5‰ and 13.0‰ (mean 12.7‰), is more variable but still very small (Table 3). Low variability in *δ*^13^C and *δ*^15^N was also observed for Verteba Cave, with slightly less enriched values for both animals and humans (36). This may reflect small differences in local isoscapes between the Dniepr and Dniester tributaries but suggests similar dietary habits for each human group concerning their relative weighting of food sources.

For Kosenivka, the small differences in the isotopic signatures do not indicate the presence of additional individuals that are as yet unidentified. According to Hyland et al. [135], intra-individual differences in *δ*^15^N of up to 1.5‰ are acceptable as coming from the same individual, especially among samples from different skeletal elements and parts of the bone. This would support the assumption that different samples represent the same individual, e.g., the maxilla and the frontal bone could both belong to individual 6.

Due to the enrichment in heavier isotope variants by trophic level, the mean *δ*^13^C and *δ*^15^N of the measured animal bones is 2.6‰ and 2.7‰ higher, respectively, than that of the measured plants (Fig 11). The offsets between the measured animals and humans are only 1‰ for *δ*^13^C and 2.9‰ for *δ*^15^N, and hence considerably lower than the expected diet–consumer offsets of 4.8‰ and 5‰, respectively. The distances between the measured humans and cereals (*δ*^13^C 3.6‰, *δ*^15^N 5.7‰) are much closer to the model offsets.

### The dietary significance of plants and animals in human consumption

In the model output, the importance of cereals (C) in human (h) nutrition is correspondingly dominant, with a share of 92% ± 0.06% (Fig 12, model 1). Higher human *δ*^15^N values correlate with a higher portion of animal protein in the diet, of up to 15% ± 0.11%, in the last years of life of individual 6 (Fig 12, model 2-3; see S3 Tables).

**Fig 12.**
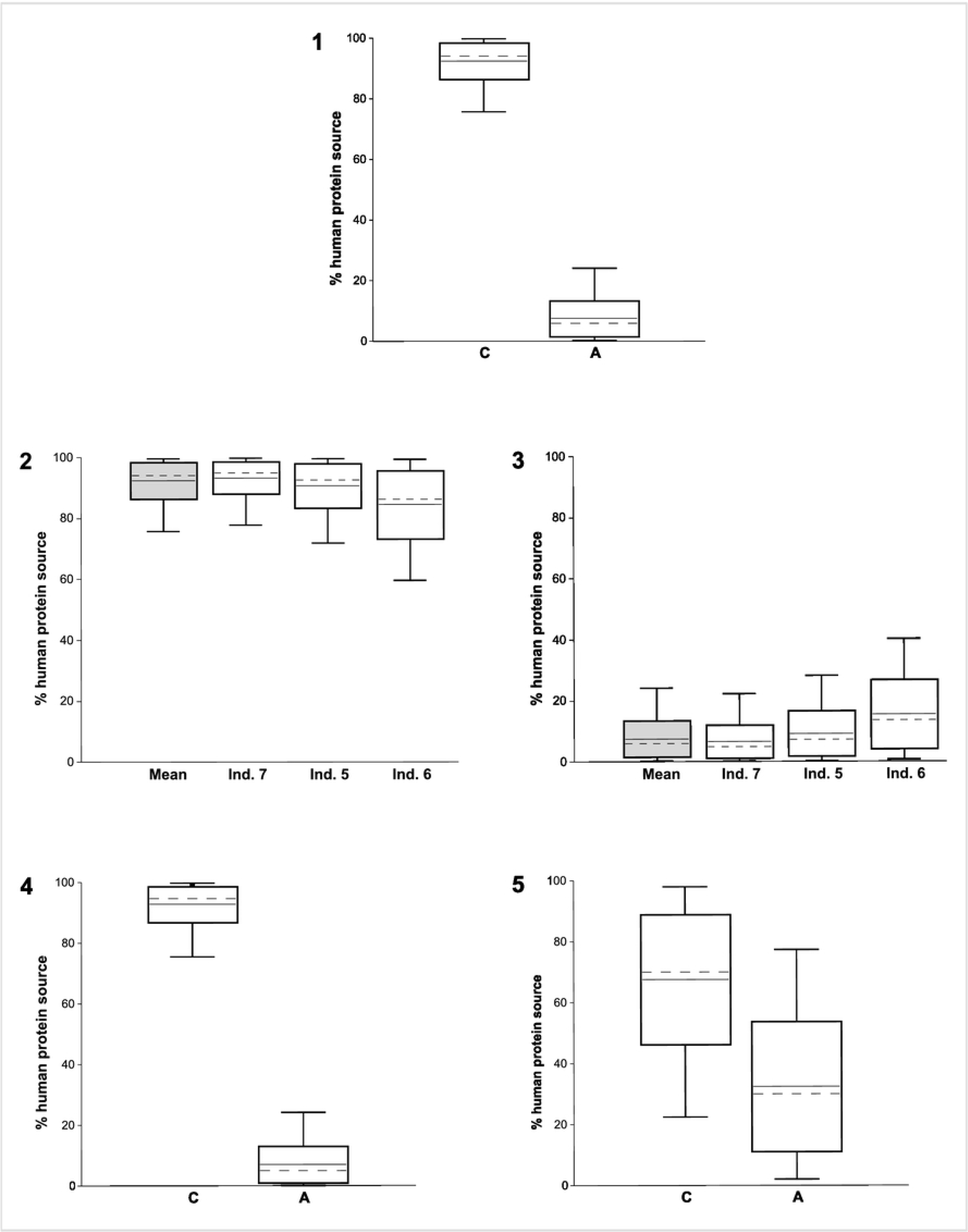
Five models of human dietary protein resources at Kosenivka using the software FRUITS, v. 3.1. [100]. (1) Proportions of cereal grain protein (C) and animal meat (A) in the diet of humans from Kosenivka reconstructed from mean δ^13^C and δ^15^N values and SEM (Table 4). (2) Proportion of cereals calculated for the human mean (h, n=3) as in model 1 and for individuals 5–7, arranged by decreasing δ^15^N in cranial bone collagen. (3) As model 2 but for the proportion of animal meat. (4, 5) Results for food source inputs to the human diet considering either δ^13^C (4) or δ^15^N (5) as a single fraction in the FRUITS model (see S3 Tables). Boxes span 1 standard deviation from the mean (horizontal line); whiskers span 2 SD; medians are marked by discontinuous lines. Illustration: F. Schlütz, K. Fuchs.

Due to metabolic processes, the calculated portions reflect primarily the protein part of the food consumed. Some 75% of the collagen carbon and almost all of the collagen nitrogen derives from food proteins [162]. Compared with meat, cereals are quite low in proteins and mostly consist of nitrogen-free hydrocarbons, such as starch. Therefore, the caloric importance of cereals, through their hydrocarbons, for human diet must have been greater than the mainly protein-based calculations suggest.

Furthermore, these estimations are based on a model restricted to the food sources cereals and meat from terrestrial animals, whose sample material stems from a domestic context. No finds or isotopic data for gathered wild plants or fish are available for Kosenivka. It seems plausible that the real diet was more diverse and included fish and collected wild fruits, as is known from other Trypillia sites [14,136]. Bivalves; secondary products of (wild) animals (eggs, milk); legumes; and vegetables are also rarely if ever represented among archaeozoological and archaeobotanical finds [137,138], and therefore their dietary significance is unknown.

To test the general role of food sources for the isotopic composition, we re-ran the model with only *δ*^15^N and again with only *δ*^13^C. With *δ*^15^N as the only fraction in the model, cereals account for 67.5% of the diet (Fig 12, model 4), and with *δ*^13^C as the only fraction, they account for 92.9% (Fig 12, model 5). This underlines the clear predominance of cereals as the source of protein in the human diet. Beside cereals, other food sources with low *δ*^13^C, such as wild plants, fruits, and vegetables, may also have played a role in the human diet. Fish bone collagen can exhibit *δ*^15^N values that match those of mammals [134] and therefore could hold the same position as meat in a food web model. Conversely, fish bone collagen can also exhibit *δ*^15^N values as low as 7‰ [139]. Without concrete finds from the site, the contribution of freshwater resources remains unclear. Hence, for more differentiated insights into the human diet at Kosenivka, isotopic signatures of additional possible food sources are required.

### The palaeolandscape of Kosenivka

To gain information on the palaeoecology of the landscape in which the settlement of Kosenivka was situated, we converted the animal bone collagen signatures to an approximated terrestrial vegetation signal by subtracting the offsets defined for the trophic stage in the food web (*δ*^13^C 4.8‰, *δ*^15^N 5.5‰). Because all the animal *δ*^15^N values from the site fall within or even below the *δ*^15^N range of the identified herbivores, the converted signatures of all animals reflect mainly the mean isotopic composition of the consumed vegetation. This may have been a mixture composed of grasses, fruits, leaves, or branches of trees and shrubs from different habitats. Drucker et al. [104] calculated the *δ*^13^C values of consumed plants based on bone collagen from recent animals that fed in deciduous forest and forest–steppe ecotones, which we used to illustrate possible levels of habitat openness for the Kosenivka dataset (Fig 13). Because the values reflect mostly the composition of vegetative plant parts, as these predominate in the animals’ diet, we adjusted the *δ*^13^C values from the cereal grains to enable us to compare the growing conditions of the plants (95,96). The isotopic values of cereal grains, individually or as a mean, stand for nutrient and water availability on the arable soils. Due to the short grain-filling season, the *δ*^13^C values reflect water availability and relative humidity in summer, a few weeks before the harvest.

**Fig 13.**
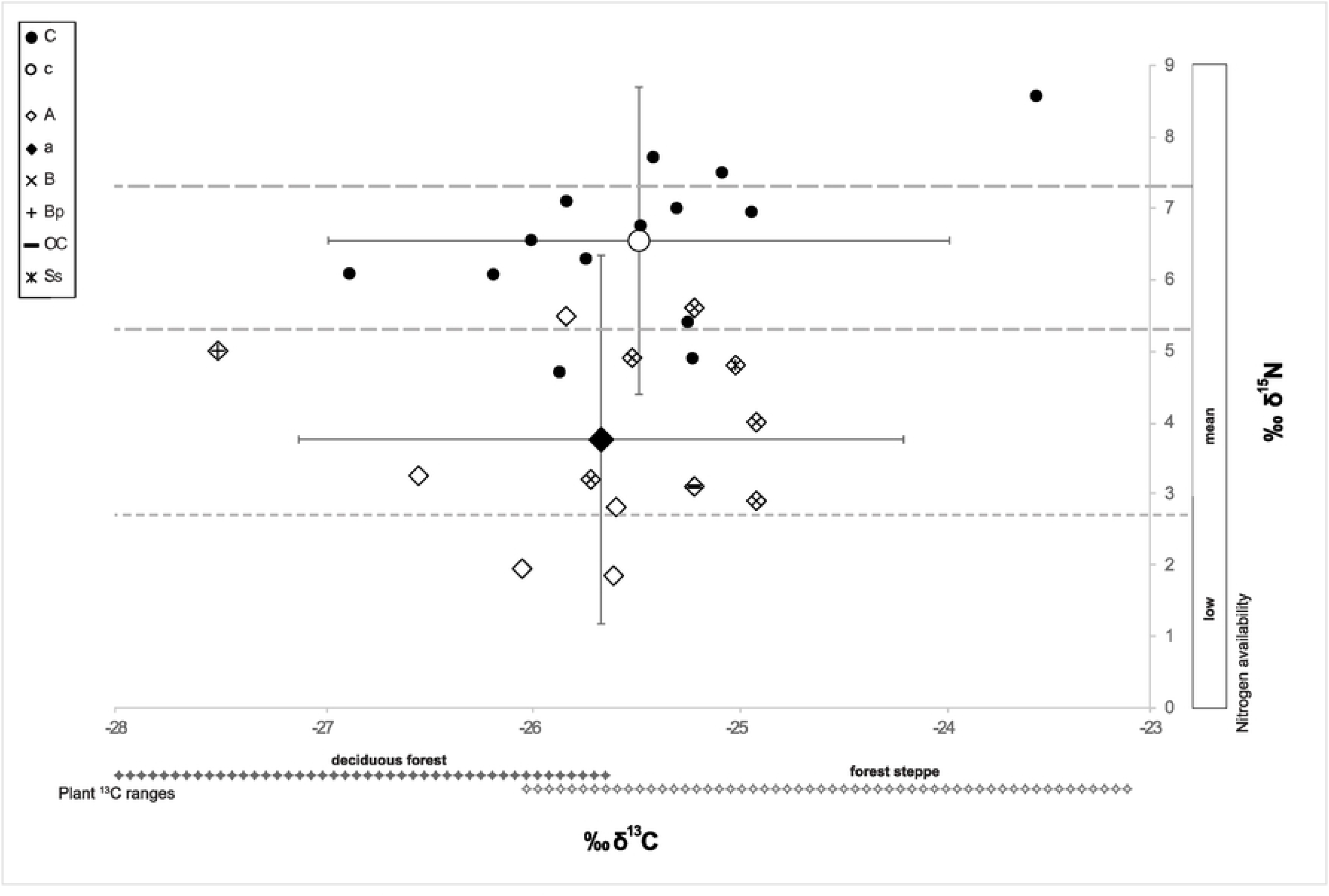
Reconstructed δ^13^C and δ^15^N isoscape of Kosenivka. δ^13^C of cereal grains adjusted by −2‰ to stand for vegetative plant parts. Values from animal bones reduced by diet– collagen offsets (δ^13^C 4.8‰, δ^15^N 5.5‰) to reflect the mean isotopic composition of the consumed plant material. Ranges of δ^13^C of consumed plant material estimated from bone collagen of recent herbivores for deciduous forests and forest-steppe from Drucker et al. [104], adapted to differences in offsets and Chalcolithic atmospheric δ^13^C concentration (see Materials and methods). Illustration: F. Schlütz, K. Fuchs.

The *δ*^13^C values calculated from the undetermined animals fall into the terrestrial plant range of the presumed transition area from forest-steppe to deciduous forest (Fig 13; [79]) Some specimens exhibit the lowest *δ*^15^N values of our data set, at around 2‰, a value often found in grasses, herbs, and tree foliage growing in forests [140,141]. In contrast to all other animals, the aurochs probably grazed in quite closed forests, where ground plants are depleted in *δ*^13^C by the canopy effect [104]. The *δ*^15^N of the aurochs diet points to a vegetation well supplied with nutrients, possibly growing in floodplain forests. Such wooded and humid habitats are typical for *B. primigenius* [142]. With *δ*^13^C values mostly higher than the overall animal mean, most of the other *Bos* specimens (most probably cattle) cluster in the forest-steppe range, with *δ*^15^N values indicating mostly good or even high nutrient supply.

High *δ*^13^C values, of around −25‰, could locate the feeding of some cattle, the wild boar, and the sheep/goat in a rather open forest-steppe or even open steppe–like environments (Fig 13). Isotopes from tooth enamel could help identify animal movements between more open and closed habitats [143,144], but animal teeth were not found at Kosenivka. The mean *δ*^13^C of the vegetative cereal parts exceeds that of the animals’ food plants by just 0.2‰. Bright, open growing conditions are manifested by high *δ*^13^C values in about half of the single measurements. The specimens with low *δ*^13^C may point to shady growing conditions under trees, locally wet soils, or simply a few rainy weeks before the harvest.

The cereals grew mostly on soils with good to high nitrogen availability. Because *δ*^15^N values of up to 5‰ may represent natural conditions in the area [145] , *δ*^15^N values in excess of 5‰ may be evidence for soil fertilisation. In the literature, values higher 3‰ are accepted to be indicative for medium levels of manuring and from 5.8‰ on for high levels of manuring [89,146]. By these cut-offs, 80% of the sampled cereal grains from Kosenivka would have been cultivated under soil conditions typical for high manuring. Manuring could have been practised by the farmers using livestock dung collected from the landscape or from kraals. Possible other reasons for the high *δ*^15^N of the cereals are some kind of field rotation [147] or the use of previously intensively pastured land as sowing areas. Intensive pasturing leading to an increase in *δ*^15^N in animals and plants was demonstrated for Trypillia sites by Makarewicz et al. [13,14]. The highest cereal *δ*^15^N values may result from locally increased nitrogen enrichment from faeces and/or urine during grazing of stubble fields or from intensive manuring. Plant production boosted by high-nitrogen soils may have led to a relative shortage in water availability in the lush-growing cereal plants, which could explain the enriched values of both isotopes in the cereal grains.

### Archaeological analogies

In previous compilations of CTS human remains, the striking contrast between the high number of settlements, often inhabited by large populations, and the extreme rarity of human remains has been highlighted as characteristic for the CTS phenomenon [20,148].

For the period between 4800 and 3000 BC, remains of a total of approximately 1058 individuals have been found (S5 Tables); of these, 1041 were recovered from distinct settlement and cemetery contexts, mostly of the later phase. This small number contrasts with the huge number of settlements over a period of 1800 years. It is estimated that there were at least 1500, some of which had an extreme size of more than 100 ha and a population size in the thousands. Where estimates of the population size of settlements are available, they mostly refer to individual sites or to the catchment area of the Sinyukha River, the region with the highest level of population concentration in large mega-sites. According to estimates by Ohlrau [23], 3500 to 4000 households existed here simultaneously in the period 3950– 3700 BCE. Based on a commonly used size of a nuclear family, given as 4.5 to 7 people per household (according to Cook; [138]), this number of households corresponds to ≈15,750– 28,000 individuals in this region [23]. Only 19% (n=193) of the recorded skeletal remains date from before 3650 BCE (Figs 14.A and 15), i.e., pre-dating and coinciding with the existence of large, aggregated settlements and peak population sizes (Trypillia A–C1), and only 8 of these 193 skeletal finds originate from the Sinyukha region. The other 81% (n=848) stem from the period after 3650 BCE (Trypillia C2; Fig 14.A), thus from a much shorter time span. Given these proportions and considering the estimated population sizes for the Sinyukha region, we can state that only ca. 0.03–0.05% of the Trypillia population is represented in the human bone finds.

**Fig 14.**
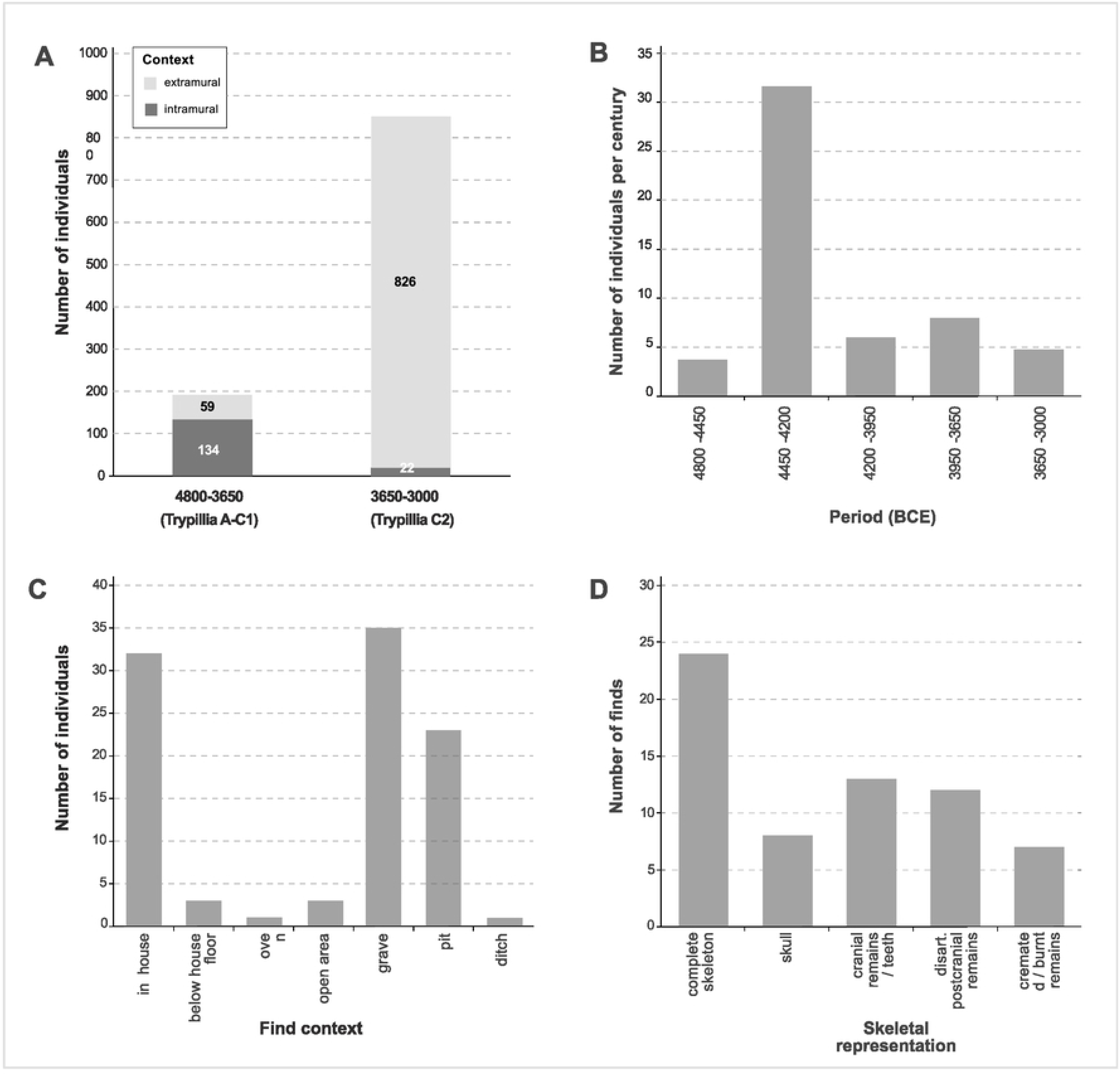
Records of human remains from Cucuteni–Trypillia contexts. (A) Frequency of human remains in intramural and extramural contexts across time (n=1041). (B) Relative frequency of human remains in settlements across time (n=163). (C) Frequency of individuals found in each category of find context in settlements (n=163). (D) Frequency of representation types of human skeletal finds in settlements (disart.=disarticulated; n=64). Illustration: R. Hofmann, K. Fuchs.

In the phase before 3650 BCE, 70% of human remains were recovered from settlements (n=134; Figs 12.A and 13) and only a relatively small proportion (30%) from extramural burials (n=59), such as at Chapaevka (Trypillia B2 or C1, 31 individuals) and Ostrog Zeman (Trypillia B2/Malice, 18 individuals). After 3650 BCE, this trend reversed, coinciding with the abandonment of the large settlements, a partial shift in geographical focus to the North Pontic Steppe zone, and the formation of the Late Trypillia groups. There is a general increase in the number of human remains from individual sites, as well as a considerable increase in the proportion found in extramural contexts (n=826, or 97%) vs settlement contexts (n=22, or 3%). Although this trend reversal is certainly due to the general decline in settlement evidence, the increase of extramural finds is confirmed by, among other things, larger cemeteries at sites of the Late Trypillia groups, such as those at Vichvatinci (MNI=72), Usatovo (MNI=88), and Majaki (MNI=38), and at some sites of the Sofievska local groups, such as Sofievska (MNI=145), Krasniy Chutor (MNI=195), and Cherin (MNI=95).

The highest density of human remains from settlements, averaging >30 individuals per 100 years, is recorded for the period 4450–4200 BC mainly for the western CTS (Figs 14.B and Fig 15, S5 Tables). Most of these human remains were found in almost equal proportions in house contexts, intramural graves, and pits, and very few were found in open spaces, in ditches, next to ovens, or below house floors (Fig 14.C, S5 Tables). It often remains unclear if below-floor finds are chronologically related to the house or pre-date it. There is one case from that period similar to Kosenivka. At the Cucuteni settlement site of Poduri-Dealul Ghindaru (Trypillia A1, ca. 4550–4300 BCE; see Fig 1) the skeletal remains of five individuals, all assumed to be children, in crouched position and exhibiting different orientations, were found in a burnt house context. Unfortunately, there is no detailed information addressing taphonomic or depositional processes. This is one example for which the researchers suggests a deathly fire event as a possible explanation; the other explanation they suggest is human sacrifice [110,151]. In the later cemeteries, inhumations in pits predominate, but cremation remains in urns occur as well [115]. Overall, and especially for settlement depositions, the diversity in skeletal representativeness and preservation is striking, ranging from complete skeletons to disarticulated parts of the postcranium, crania, and cranial fragments (e.g. [20,88; 191–212, 251–254]; Fig 14.D, S5 Tables). Some of the human remains from settlements show fire impact, ranging from charring to complete calcification, and these cases are not restricted to burnt house contexts (Fig 14.D).

**Fig 15.**
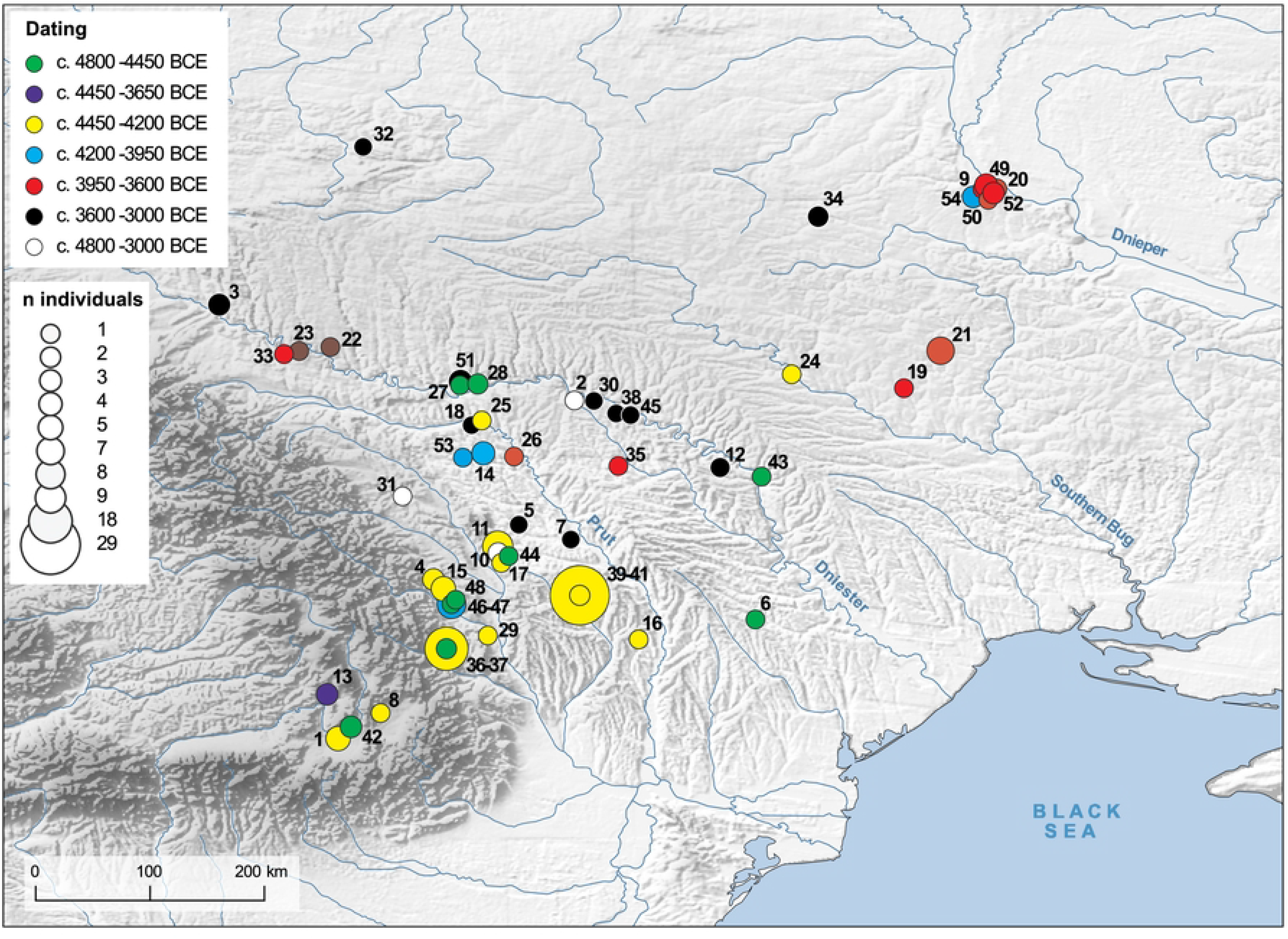
Spatial distribution and dating of human remains in CTS settlements. The numbering corresponds to the site identifiers in S5 Tables. Illustration: R. Hofmann.

In this review of the complex record of CTS human remains, the Kosenivka finds stand out because they mark the transformation period during which the change from predominantly intramural to predominantly extramural depositions took place (Fig 14.A). Because researchers have observed such great diversity among CTS human assemblages and because the numbers demonstrate a striking underrepresentation of human remains compared with estimated population sizes, data from other sites yield few clues to how the finds from Kosenivka are to be interpreted. We have to consider the possibility of either a lethal fire event, as suspected by the excavators; a violent death; or more complex formation processes involving ritual practices.

## Discussion

### Insights into Trypillia life

#### Subsistence and diet

Knowledge on the food economy in the early history of the Bug–Dnieper region is very limited [152,153], and isotopic data for CTS sites are mostly available only for faunal remains. Until now, Verteba Cave was the only site with isotopic data on faunal and human remains, but it lacks the domestic link [36]. Our knowledge on the CTS plant economy has recently been improved through implementation of systematic sampling, careful bucket flotation, and “pit archaeology”. The focus on house-accompanying pits, which have been shown to be the best context for the recovery of plant and animal remains, allows an insight into the spectrum of foods consumed and its production in a household [14].

Thus, to date Kosenivka is the first Trypillia site for which stable isotopes from human, animal, and cereal remains enabled detailed insights into Chalcolithic subsistence. Despite the overwhelming dominance of animal bones in the archaeological find assemblages from other settlement sites, our food web calculation clearly unmasked plants as the primary food source for the Trypillia people at Kosenivka, making up some 90% of the daily protein nutritional portion. Proteins represent roughly less than 20% of the caloric content of cereals. Carbohydrates (sugar, starch) and a small amount of fat together contribute some 80% of the energy, but neither of these energy sources contain nitrogen, and they contribute only a small proportion of the carbon synthesised in bone collagen [103]. Consequently, cereal grains and any plants of related composition are *a priori* underrepresented as a source in calculations in which consumers are represented by their collagen. The true caloric contribution of plants to Kosenivka human nutrition must be assumed to have been higher than the modelled 90%.

The remaining proportion, representing the level of meat consumption of the Kosenivka inhabitants, may have been quite close to the lowest levels reported for Neolithic and for historic farming communities, representing just a few kilograms per year [157]. Based on this, we can assume that the production of manure was a much more important part of animal husbandry than the production of meat. Even today, in farming societies without access to artificial fertiliser [158], large dung heaps are considered symbols of wealth, as they mark how many fields a household can fertilise [159]. Furthermore, our results support the hypothesis that cattle were prestigious animals, primarily used as draught animals to farm the fields, for dung production, and for dairy product production, rather than for meat production. We note that the role of pigs at Kosenivka remains unclear in this respect, as we lack isotopic data. At Verteba Cave, the small isotopic distance in *δ*^15^N between human and pig bone collagen is thought by Lillie and colleagues [36] to suggests that pork was not an important protein source for the humans. But because the bones were found in a non-domestic setting, it is not certain that there is a direct dietary connection between these faunal and human values. The authors of a recent publication presenting isotopic values of 12.0‰ *δ*^15^N and −20.1‰ *δ*^13^C obtained from a human female long bone found at the Trypillia settlement of Kolomiytsiv Yar Tract (4049–3820 cal BCE, 2σ range, ca. 150 km north of Kosenivka) assume that the values reflect either a forager–pastoralist diet or a forager–pastoralist diet with an admixture of arable crop products for this individual [160]. The authors suggest that the enriched nitrogen values, of 11.5–13‰, indicate consumption of some freshwater resources. But according to our results, it is possible that at Kolomiytsiv Yar Tract the high nitrogen values are due to a cereal-dominated diet, just as they are argued to be at Kosenivka. Thus, our results give new food for thought on interpreting CTS isotopic values that fall outside the range of Eneolithic farmers on the West Pontic Coast, of Eneolithic forager– pastoralists on the lower Dnieper, and of Eneolithic to Early Bronze Age pastoralists on the Pontic Steppe, according to groups classified by Nikitin and colleagues [160, see Fig. 3] .

We need additional isotopic information to (i) better understand the contribution of different plants and animals to the human diet, such as δ^13^C from bone or tooth apatite [161,162]; (ii) provide a more differentiated picture of the human diet, including δ^13^C and *δ*^15^N from additional food sources, such as collected plants; and (iii) increase the number of input sources in order to reduce uncertainties in models of animal and human diet, such as δ^34^S [152,163,164]. Nevertheless, our major result highlights the extremely great importance of cereals in the diet of the Kosenivka individuals. This is consistent with the indication of manuring given by the high nitrogen values of the grains. To supply the large mega-site communities with the cereals they would have required, manuring of the arable fields was necessary to ensure an adequate harvest, and the production of dung may have been the main motive for animal husbandry. Thus, our reconstruction of the food web assigns a different importance to livestock in the food economy of the Trypillia mega-sites than it had previously been assigned.

#### Oral health, disease, and living conditions

Oral health is heavily influenced by the composition and processing of consumed foodstuffs. Observations made for the maxilla from Kosenivka and its four preserved teeth provide further information in this regard. The isotope results demonstrate the large dietary role of einkorn and emmer, which is supported by the molar dental wear induced by a highly erosive food substance. Because such shallow dental wear patterns are associated with agriculturalists, they emphasise the role of cereal products in the diet of the people whose remains were excavated at Kosenivka. Interestingly, the four preserved teeth were free of caries, a finding that would not be expected in an adult of advanced age who had consumed large amounts of cariogenic carbohydrates. However, the aetiology of caries is multifactorial, including individual (genetics; enamel mineralisation, saliva components, hormonal state), environmental (flour concentrations in ingested water, diversity, and abundance of oral microbiota), and dietary (stickiness, starch and sugar contents) factors [57,168,169]. In addition, highly abrasive food is suggested to prevent occlusal caries [170]. Dental calculus of considerable accumulation, as observed on all studied maxillary teeth, results from an alkaline pH oral milieu, which is associated with a protein-rich diet and a lack of dental hygiene [171]. Interproximal grooving suggests the occasional consumption of fibrous food that tends to stick in the interdental spaces [172], which could be associated with meat or fish; but has also been observed scraping carious lesions [125]. An occupational causation of these grooves is less likely, as occupational grooves are more often associated with a notch-like appearance rather than sharp horizontal striations [173], but this is not mutually exclusive [174]. However, because the described groove is very faint, the cause remains unclear. The amount of observed enamel cracks or “chipping” results from biting and chewing substances harder than the tooth itself. This occurs when food components include firm substances, e.g., impurities from cereal processing or shell from cracking nuts, or when the individual uses their teeth in other occupational activities [126,174]. Thus, the sparse oral health data of this individual are in line with a mixed economy including animal protein and cereal processing but do not give decisive information on eating habits of the wider population.

Despite the poor skeletal representativeness, several pathological markers have been observed on the cranial remains of the Kosenivka individuals. Understanding the causes of unspecific skeletal phenotypes is a major concern in the field of palaeopathology. The causes cannot even be reduced to biological stress, as we know from clinical research that psychosocial stress triggers high cortisol levels, which can manifest in inflammatory processes (e.g., [143]) Yet, pathological markers have been observed on 7 of ca. 20 assessable cranial bones. This proves the presence of stressors in the life of the Kosenivka inhabitants. For the periosteal changes in the oral cavity, in the maxillary sinus, and on the endocranial surfaces, infections with bacteria or viruses are likely causations [58,118].

For the Dnieper region, palaeopathologial analyses of prehistoric skeletal remains have been published by Lillie and colleagues [176], Potekhina [177] and Karsten et al. [178]. Physical stressors and oral health data are used as proxies to trace the biological consequences of foraging and farming subsistence strategies and the transition between the two. According to the Lille et al. [176] data set, based on 189 skeletons spanning the Epipalaeolithic to the Chalcolithic, *cribra orbitalia* was observed once in a Neolithic individual (Dereivka I). In line with low frequencies of dental hypoplasia (1.9% of teeth), this tempted the authors to conclude on low levels of physical stress, which would be in contrast to the individuals found at the 350 km distant Verteba Cave (18.2% of teeth; [144]). The adult frontal bone from Kosenivka (individual 5/6/+, constitutes the second published case of *cribra orbitalia* in a CTS assemblage, proving that biological stress reflected as an increased demand for red blood cells was an issue in Trypillia times too). Enamel hypoplasia, whose solely stress-caused aetiology has recently been challenged [179], was not observed on the three teeth. The advanced dental wear is a limiting factor in establishing the presence of enamel hypoplasia, especially for those defects developed early in life. The absence of caries in the Kosenivka individual fits with the overall low incidence of caries in the Dnieper Mesolithic, Neolithic and Chalcolithic populations, and the same applies to the accumulation of dental calculus [176]. There is one other CTS case of “tooth-pick grooves”, reported for an individual from the Neolithic cemetery of Vovnigi II (5470–4750 cal BC, Dniepr River rapids) and interpreted as being the result of an oral hygiene habit aimed at removing dental calculus. Unfortunately, there is no detailed information for direct comparison

Potekhina [177] sees the decline in life expectancy from Neolithic fisher-hunter-foragers (29.5 years) to the fully agriculturally established Late Trypillia populations (20.9 years) as having been caused by their subsistence strategies and the consequences for health of the farming lifeway [176]. Unfortunately, Kosenivka does not offer much statistical potential in this respect due to the imprecise age-at-death estimations, especially of the adult individuals. The types of pathological markers observed for the Kosenivka individuals are in line with other Neolithic populations (e.g., [126–128]). Unfortunately, their number is too small to allow for further, supra-regional interpretations. At this point, we cannot infer a relationship between disease burden and changes in settlement patterns during this late phase of the CTS. It remains to be investigated whether the possibility of achieving a reduction in the negative health effects of living in mega-sites may have been a reason for the shift to a more dispersed settlement pattern.

#### Traumatic injuries: Interpersonal violence?

Perimortem cranial trauma most probably suggest that two adults among the Kosenivka individuals received interpersonal violence and less probably that they received trauma of accidental causation. The postcranial elements do not show traumatic injuries, but this does not mean such injuries were not present originally. Better preservation of the postcranial remains would have helped us to evaluate the potential for conflict and the risk of acquiring injuries for the Kosenivka inhabitants. Yet, two of four larger skull fragments that could be tied to an individual are affected, which is noteworthy because up till now, the Verteba Cave cases were the only evidence for interpersonal violence in Trypillia times.

The Middle Trypillia finds from Verteba Cave have been described as being unique for providing the most direct evidence of conflict for the CTS cultural phenomenon related to the Trypillia B2/C1 period [183]. The assemblage represents a minimum of 36 individuals of different biological sex and age-at-death, 25 of whom are represented solely by cranial remains [184], in some cases deposited in “skull nests”. Of these cranial remains, 11 exhibit several signs of cranial trauma, with the majority showing vault-penetrating trauma (with a lesion size up to 63 mm) that proved lethal, although two survived for some time after being injured. Most of the individuals had been hit on the back of the head [184], and stone or copper tools are suggested as weapons [183]. In addition, the bones showed cut marks indicating severing of the head from the vertebral column [183,184]. Based on the information available, we can state that the shape and locality of the injury of Kosenivka individual 1 is similar to the depression fractures observed on the Verteba Cave specimens. Our review of the literature found evidence for acts of interpersonal violence post-dating Trypillia stage C1, that is, after the decline of the mega-sites and the shift towards burials in cemeteries, and thus occurring in a differently organised society. Publications on the human remains recovered from the cemetery of Majaki, which is associated with the Late Trypillian Usatovo, describe traumatic injuries that are apparent on the crania of two adult males (mound 1, burial 9; complex 3, burial 2a) out of a total of 38 individuals [185,186]. One of these males exhibits six injuries, on the frontal, parietal, and occipital bones. At least one trauma is indicative of a lethal course because the lesion penetrated the vault without signs of healing. From the lesion shapes and fraction patterns, the authors suspect a stone axe as the weapon that inflicted the trauma [155, Figs 1-2, 4].

The record of perimortem trauma in CTS is strongly biased: for the earlier Trypillia periods, especially before late C1, we lack reliable figures on the number of skeletal remains and thus reliable figures on perimortem trauma. It is noteworthy that the individuals with perimortem cranial trauma are from very different depositional contexts: a cave, a house, and a cemetery. Motivations for the violence remain unclear, and these may include socio-political or economic stressors, as recently discussed for the Neolithic in northwestern Europe [187].

Interpretation of the exceptional deposit of selected and severed skulls showing evidence of violence in the Verteba gypsum cave system is the subject of controversy. Madden and colleagues [184] see the traumatic injuries of the Verteba Cave individuals as proof that interpersonal violence was part of later Trypillia times, perhaps connected to subsequent changes in cultural dynamics in the Danubian Chalcolithic. Others [116] have proposed a multicomponent, ritual funeral process for the Verteba Cave rather violence (similar to the process argued for some Linear Pottery mass grave sites; [172,173]). A ritual rather than a conflict dimension of violence is also possible for the domestic sphere, including at Kosenivka, although here, whether there is a link between the possibly lethal cranial trauma and the scenarios described below for house 6 remains can be questioned. Kosenivka house 6 and Verteba Cave (both C1) prompt us to discuss how violence was handled in the cooperative societal systems that are assumed for the mega-site phase of the CTS [190].

### Scenario for Kosenivka house 6

One of the central aims of this paper was to scrutinise the hypothesis of the discoverers that the human remains in house 6 represent individuals who died in a fire event and to discuss other possibilities. To address this hypothesis, we generated more information on the number of individuals, their demographic profile, and signs of violence on their bones; assessed bone alterations due to fire impact and taphonomic processes; and refined the chronological models by further ^14^C dating.

The bone assemblage is characterised by a high degree of fragmentation, low skeletal representativeness per individual, and differential impact of fire. The fragmentary status of the skeletal elements and the vast dispersal across the archaeological structures is likely due to the near-surface location of the entire feature and to post-depositional damage, probably erosion and modern agricultural activity. Information on the *in situ* situation of the isolated bones is sparse, but re-fits between bone fragments from different excavation squares prove that at least some bones from a single individual were dispersed several metres.

In general, our osteological re-examination resembles the original results by Kruts et al. [21] although it increases the MNI from 4 to 7. We were able to obtain data that supports the authors’ assumption that these individuals represent a broad age-at-death distribution (children, adolescents, younger and middle-aged adult individuals). Although the osteological markers are sparse due to poor preservation, we suggest that the assemblage contains females as well as at least one male. There is no evidence of selection in terms of age-at-death or biological sex; rather, the estimated biological profile matches the expected composition of a household, perhaps a family.

Burning, with combustion temperatures of 550–800°C, impacted only the bones found inside the central house structure. Recent experimental research on pieces of daub demonstrates combustion temperatures of 750–850°C for a domestic structure and 650–750°C for a communal building at the Trypillian mega-site of Maidanetske [191]. Most of the burnt human bones from house 6 at Kosenivka show evidence for combustion at temperatures within the ranges demonstrated for Maidanetske, and the nature of the burnt assemblage is very atypical for intentional cremation [72]. Our macroscopic and histotaphonomic analyses on the burnt and unburnt remains provide some evidence that combustion took place when the bones were still fresh, and that the time between the death of the individuals and the fire was probably not longer than a few months, as evidence for the diagenetic breakdown of collagen structures by microbes as part of natural decay was strong in the unburnt specimens but completely absent in the burnt specimens. Two further aspects are important for drafting scenarios. Firstly, the unburnt assemblage consists of cranial and long bone fragments only, whereas the burnt assemblage shows greater skeletal representativeness. Secondly, the individuals showing perimortem cranial trauma represent both assemblage groups: burnt from within the house structure (individual 1) and unburnt from the outer activity zone (individual 5).

The ambiguity around several aspects of the taphonomic processes has hampered us in deriving possible scenarios. The first aspect is the occurrence of only unburnt skull and long bones in the house periphery. It is still unclear if this is due to preservation or selective, probably secondary, deposition. Bones of the hands, feet, pelvic girdle, and shoulder girdle, as well as the rib cage, are prone to be biased against in the archaeological record due to their size, shape, and structure [192]. Yet this same biased representation could argue for the skull and long bones having been selected as part of a multi-phased funeral activity. The second aspect is the lack of gnawing marks. It is still unclear whether this lack shows that the human remains were not accessible to scavenging animals, e.g., by being covered by house debris or soil; that scavengers were kept away from the house; or that scavengers were absent. The third aspect is the lack of traceable microbial damage in the burnt assemblage. The abundance, type, and activity of bacteria and fungi are decisive for the onset and extent of microbial bone alteration. Influential factors are post-mortem treatment of the corpse, such as dismemberment or defleshing, and if and in what microenvironment it is buried – not only compactness, humidity, oxygen availability, and pH, but also the individual’s microbiome composition [59,65,68]. Thus, exactly determining the time interval between the death of individuals 1–4 and the combustion of their physical remains is impossible because it is unclear whether, when, and how they were covered by soil. However, results from experimental studies indicate that an interval of between days to three months [64,193] between these two events.

The probably lethal cranial trauma of individuals 1 and 5 is also of importance in this regard. On the one hand, this trauma could be evidence for a connection between a violent act and an intentional fire and on the other hand, it could indicate that their deaths and the fire were two completely different events.

The refined radiocarbon chronology obtained from both unburnt and burnt human bones, as well as unburnt animal bones, for house 6 suggests an extremely short period of deposition for all but one of the bones, of less than 35 years (the exception is a bone deposited approximately 120 years after the house burnt down, on top of the remains of the house). The dates indicate at least two and more likely three depositional events.

The first took place before 3900 BC and involved the deposition of animal bones representing waste from the time of house use. The deposition of human bones representing the inhabitants of the house at the time of the fire event was either part of that deposition or a separate, second, deposition. After the inhabitants of house 6 died, house 3 was probably also abandoned. The second or possibly third depositional event represented by the material in our study is the deposition of a human skull on the rubble of house 6, several generations after the house burnt down.

An alternate scenario, i.e., that the human remains represent a *rite de passage*, is less probable in our option, because of the short duration of use of house 6; the lack of comparable finds from the other four excavated houses at Kosenivka and from the entire region; the taphonomic observations, especially the lack of gnawing marks; and the time that elapsed between these individuals’ deaths and the fire event, which was insufficient for microbes to feed on bone tissues and for complete dehydration of bone organic compounds. The most probable scenario is therefore a single depositional event, with a possible connection between fire and death. Coinciding with the hypothesis by Kruts and colleagues [21], in this scenario, members of a household died during a single event, e.g., a fire event that was more devastating within the domestic structure than in its surrounding activity zone. Individuals 1–4 died in the house and their remains combusted inside the house, whereas individuals 5–7 may have managed to escape outside and perhaps died by carbon dioxide poisoning, their skeletal remains left unburnt. It remains unclear if this fire was accidental or an act of violence, although the cranial trauma of individuals 1 and 5 would support the latter. In this scenario, the house was not inhabited after the death of the individuals.

Different from this is the deposition of the skull from individual 7 around 3500 cal BC (dating to stage C2), which is surely not the result of an accidental fire that killed a whole family, but of a deliberate ritual deposition.

### Trypillia human remains

Our literature review of the published record of CTS human remains from intramural and extramural contexts has demonstrated that (i) most of the finds date to the Late Trypillia phase, especially after ca. 3650 BCE;

(ii) there is great diversity in the type of depositional context and the condition of human bones from intramural contexts; and (iii) there is a transition between stage C1 and C2 from rare, intramural, mostly disarticulated and isolated bone finds, to predominantly extramural burials. The studied individuals from Kosenivka add to the ca. 0.05% of the CTS population of the Middle Trypillia phase that are represented by skeletal remains in the Sinyukha region before 3650 BCE.

However, the question remains: Could some of these intramural finds of human bones represent an element of a *rite de passage* that involved keeping human bones inside houses, rather than represent the remains of random events?

The observed diversity among intramural human depositions may be the result of complex, multi-stage mortuary sequences that include post-mortem rules and practices for dealing with deceased individuals. Related formation processes, or “deathways”, may have included secondary relocation, targeted removal, and selective accumulation of body parts, as is suggested by the collection of human skulls that was found in Verteba Cave [116]. The rarity of human remains compared with the estimated population sizes of the Trypillia populations raises the urgent question of whether the settlement finds represent a specific population segment and whether they relate to burial practices. Is this pattern the result of a “visible” part of a funerary circle or is it the outcome of extra-funerary formation processes, related to, for example, a bad death, low social status, sacrificial ritual, or the taphonomic loss of corpses [117].

In contrast to the situation in the eastern Balkan area [109,110,194], in the Cucuteni–Trypillia network, the overwhelming proportion of the populations remain invisible in the record of human remains. In that respect, CTS is very similar to other parts of southeastern Europe [148,195,196]. In line with current research discourse, this means that the majority were part of a different *rite de passage.* We lack in-depth studies on taphonomic processes – studies that would add to our understanding of how the dead were treated. In addition, it is still an open question which criteria led to the decision on which treatment a deceased received. So far, the sparse anthropological data do not reveal any suspicious overrepresentation regarding either sex or age-at-death (e.g., [89]).

## Conclusion

Our comprehensive interdisciplinary analysis of the human remains from a Trypillia house at the settlement of Kosenivka, which are among the few preserved human bones for the Trypillia mega-site phase, included bioarchaeological and archaeological information, from isotope studies to the nature of depositional processes.

The Kosenivka human bone data set is exceptional for the anthropological information we were able to obtain from it, despite the poor skeletal element representation among individuals. The assemblage allows rare insights into the life and death of Trypillia people who were inhabitants of the large settlement of Kosenivka towards the end of the mega-site phenomenon, probably individuals of a household, consisting of children, an adolescent, and adult females and males. Their bones and teeth exhibit traces of disease, biological stress, and dental hygiene practices, and the perimortem cranial trauma observed in two adults suggests they incurred interpersonal violence. The compilation of data on the localisation of the human bone fragments in the feature, their radiocarbon dating, and their macroscopic and microscopic characteristics of fire impact and taphonomic processes led to explanatory statements for the find situation in house 6, in which death by fire is the most probable scenario.

The six individuals who lived, and presumably died, contemporaneously are directly related to the duration of use of house 6, thus providing an estimate of the household size. The house area, of about 54 m² (12×4.5 m), fits surprisingly well with demographic estimates of the number of inhabitants of Trypillia houses based on ethnographic and ethnohistorical comparisons [22–24].

Kosenivka is the first CTS site for which carbon and nitrogen isotope values for both humans, animals, and plants are available, thus allowing us to calculate food webs. The results clearly indicate the overwhelming role of cereals and possibly other plants in the daily human diet.

Animal husbandry may have been an integral part of plant food production by providing dung as manure to enable the high cereal yields that would have been needed to feed the large populations inhabiting the mega-sites. A more differentiated insight into the diet of Trypillia people than we have provided here will require further isotopic explorations, i.e., *δ*^13^C and *δ*^15^N measurements from other possible food resources, such as wild plants, legumes, and additional animals; *δ*^13^C measurements on human bone and tooth apatite; or *δ^3^*^4^S to improve the food web models.

Our study on Trypillia human remains record demonstrates the significant shift in find contexts of human remains, from a few intramural burial depositions in settlements in early and middle Trypillia (4800–3650 BCE, MNI of ca. 197), representing less than 0.05% of the population, to extramural burials and cemeteries in the later phase (3650–3000 BCE, MNI of ca. 848, representing 81% of the entire CTS human remains record). In this regard, the human remains recovered from house 6 add considerably to the rare Trypillia human bone finds dating before 3600 BCE.

In conclusion, we argue the following:

- Population estimates: The number and biological profiles of the individuals who died contemporaneously at the Kosenivka residential house 6 (as indicated by the Bayesian modelling) indicate an inhabitation by a family or a household consisting of at least six members (two children, one adolescent, and three adults, representing both sexes). This is the first archaeological and anthropological evidence that supports population estimations of previous studies based on ethnographical and ethnohistorical analogies.
- Food economy: Food security at the Trypillia mega-sites was dependent on cereal production; meat and dairy products played a lesser role. Nitrogen stable isotope analyses have shown that at Kosenivka, the economic investment in animal husbandry was governed by the demand for fertiliser to ensure a high cereal yield, and this intensive form of agriculture, in turn, allowed for population agglomeration in mega-sites. In light of this finding, the parameters used for calculating the carrying capacities of arable land have to be reconsidered.
- Biological stress and disease: Despite the poor skeletal element representation per individual, the human skeletal remains from Kosenivka provide information on disease burden. We anticipate that thorough osteological, palaeopathological, and aDNA studies on other Trypillia human skeletal material will help to understand whether the socioeconomic stressors of living in a mega-site led to a higher disease burden, and whether this burden may have prompted the shift to a dispersed settlement pattern with potentially better living conditions.
- Ritual practices and violence: Including the two from Kosenivka, cases of interpersonal violence are currently known from two different Middle Trypillia contexts that are contemporary with the Trypillia mega-site phenomenon. The dimensions and roles of violence within cooperative societies have to be analysed further, rooted in better information, i.e., from more detailed osteological investigations of available skeletal remains. In this regard, especially the intramural human bone finds dating before 3600 BC need to be further investigated. Such investigations will also be relevant to the discussion of ritual practices in the context of death.

Considering the knowledge we were able to gain from the skeletal remains of so few individuals, it is clear that there is huge untapped potential in other finds of CTS human remains, even if these are rare in relation to estimated population sizes. We hope that the many unresolved issues will be pursued in future archaeological and bioarchaeological research.

## Acknowledgements

This article was written *in memoriam* to Helena O. Yakubenko, Vladimir A. Kruts, and Svetlana I. Kruts, great scientists of their time researching the Trypillia–Cucuteni phenomenon. Together with Galina M. Buzian and Alexey G. Korvin-Piotrovsky and other colleagues, they excavated and presented the site of Kosenivka in 2004, as part of their academic legacy. We owe gratitude to Nina Ses, curator at the State Historical and Cultural Reserve “Trypillia Culture” in Lehedzyne, for her cooperation and trust in providing the bone assemblage for analyses. We owe many thanks to John Meadows, Helene Rose, and Tomasz Goslar for comments on the radiocarbon section; Giacomo Bilotti for comments on the demographic aspect; and Stefan Flohr for comments on the histological study. We furthermore acknowledge Steven Bouillon and Yannick Stroobandt for the time and care they invested in the plant isotope measurements, as well as Cheryl Makarewicz and Rebecca Eckelmann for the zooarchaeological identification of the animal bones. We greatly thank Suzanne Needs-Howarth for English-language and text editing.

## Supporting information captions

**S1 Tables. Human remains.** Primary information and methodology: osteology, paleopathology, fire impact, bioerosion, histotaphonomy.

**S2 Appendix. Human remains**. Skeletal inventory and microscopy photos with descriptions.

**S3 Tables. Radiocarbon dating and stable isotopes.** Sample information, crude results, and FRUITS model data.

**S4 Appendix. Radiocarbon dating.** Sample quality, OxCal curves unmodelled and modelled data.

**S5 Tables. Archaeological analogies.** Data on Trypillia human finds from settlements.

